# Glutathione protects hematopoietic stem cells from proteotoxic stress

**DOI:** 10.64898/2026.10.05.756790

**Authors:** Animesh Paul, Harrison Tom, Salma Merchant, Julia D. Phan, Andrew Lemoff, Thomas P. Mathews, Courtney Karner, Gerta Hoxaj, Zhiyu Zhao, Sean J. Morrison

## Abstract

Glutathione is a redox buffer, best known for preventing oxidative stress and ferroptosis, but rarely studied in stem cells. We studied glutathione function in hematopoietic stem cells (HSCs) by deleting the glutamate–cysteine ligase catalytic subunit (GCLC), which synthesizes glutathione, or glutathione peroxidase 4 (GPX4), which employs glutathione to prevent ferroptosis. *GCLC* deficiency eliminated glutathione from hematopoietic cells, severely impairing HSC function and hematopoiesis. In contrast, *GPX4* deficiency did not affect HSC function and had little effect on hematopoiesis, even after vitamin E depletion, suggesting glutathione has ferroptosis-independent functions in HSCs. Consistent with this, *GCLC* deficiency increased reactive oxygen species levels in hematopoietic cells. But treatment with antioxidants or ferroptosis inhibitors gave little or no rescue of HSC function despite reducing ROS levels. Rather, *GCLC*-deficient, but not *GPX4*-deficient, HSCs exhibited unfolded protein accumulation, proteostatic stress, and cell death marked by elevated p53 levels and Annexin V staining. Deficiency for *p53* prolonged the function of *GCLC*-deficient HSCs after transplantation. Glutathione is thus necessary in HSCs for proteostasis but may be dispensable for ferroptosis inhibition in vivo.

## Introduction

Glutathione is an abundant redox buffer in mammalian cells^1^. It is synthesized in a two-step process catalyzed by glutamate-cysteine ligase (GCLC) and glutathione synthetase. There are no known alternative mechanisms for glutathione synthesis or importation into cells^2^. Glutathione provides a reservoir of reducing equivalents that can be used by enzymes, such as glutathione peroxidases (GPXs)^3^, to prevent oxidative stress or reverse oxidative damage. For example, glutathione is used by GPX4 in many cells to reduce lipid peroxides, preventing ferroptosis, a form of cell death marked by lipid peroxidation^4^. Inhibition of glutathione synthesis is sufficient to induce ferroptosis in some cells^5^, though many cells are insensitive to glutathione depletion^6–8^. The consequences of glutathione depletion in stem cells are particularly poorly understood as this has not been studied in most stem cells. Glutathione depletion does impair muscle regeneration^9^, though this can occur independent of stem cell function^10^; therefore, it is uncertain whether glutathione is necessary for stem cell function in vivo.

Ferroptosis limits the survival of some cells and can be induced by depleting cysteine or glutathione^11,12^. Ferroptosis contributes to the development of certain degenerative diseases^13–15^, undermines organ function during transplantation^16^, and limits the survival of cancer cells during metastasis^17^. Ferroptosis also limits the survival of hematopoietic stem cells (HSCs) in culture^18^ as well as in the absence of the histone deubiquitinase, MYSM1^19^, in vivo; however, *GPX4* deficiency does not deplete HSCs in vivo, undermine their capacity to reconstitute irradiated mice, or have much effect on hematopoiesis^20^. Cells with an HSC surface marker phenotype were depleted in vivo by combined *GPX4* and vitamin E deficiency, though HSC depletion was not confirmed in this study using functional assays^20^. GPX4 is necessary for T cell proliferation and terminal erythrocyte differentiation^20,21^, though the loss of *GPX4*-deficient red cells during terminal differentiation may reflect necroptosis, not ferroptosis^22^. Together, these studies raise the question of whether ferroptosis regulates normal HSC function in vivo.

Beyond regulating ferroptosis, glutathione also has other functions, such as regulating disulfide bond formation, protein folding^23,24^, proteostasis^25^, and the formation of iron-sulfur clusters^26^; however, glutathione is not required for all oxidative protein folding and glutathione depletion in some cells does not cause proteostatic stress^27^. Nonetheless, HSCs are unusually sensitive to proteostatic stress. HSCs and stem cells in other tissues exhibit lower rates of protein synthesis than restricted progenitors, even when controlling for differences in cell cycle status^28–30^. HSCs appear to limit protein synthesis to avoid the accumulation of misfolded proteins^31–33^. Unfolded protein or the integrated stress response promotes proteostasis in HSCs^34,35^ but can also promote HSC depletion^36^, such as by PERK-mediated induction of apoptosis^37^. In this study, we show that glutathione is necessary for HSC function but treatment with antioxidants and/or ferroptosis inhibitors in culture or in vivo gave little or no rescue of HSC function after *GCLC* deletion. Rather, *GCLC*-deficient HSCs exhibited proteostatic stress marked by an unfolded protein response, elevated p53 levels, and increased Annexin V staining. Glutathione is thus necessary for proteostasis in HSCs.

## Results

### Glutathione is cell-autonomously required for HSC function

To begin to investigate glutathione function in hematopoietic cells, we performed targeted metabolomic analysis on whole bone marrow cells, CD150^+^CD48^-^Lineage^-^Sca1^+^c-kit^+^ HSCs, and Lineage^-^Sca1^-^c-kit^+^ (LK) myeloid progenitors to assess the levels of reduced (GSH) and oxidized (GSSG) glutathione. The markers and flow cytometry gates used to identify each cell population characterized in this study are shown in Supplementary Figure 1. HSCs had nearly twice as much GSH per cell as whole bone marrow cells and a significantly higher GSH/GSSG ratio as compared to whole bone marrow cells and myeloid progenitors (Figure 1A-C), raising the possibility that HSCs may be particularly dependent upon glutathione.

**Figure 1.**
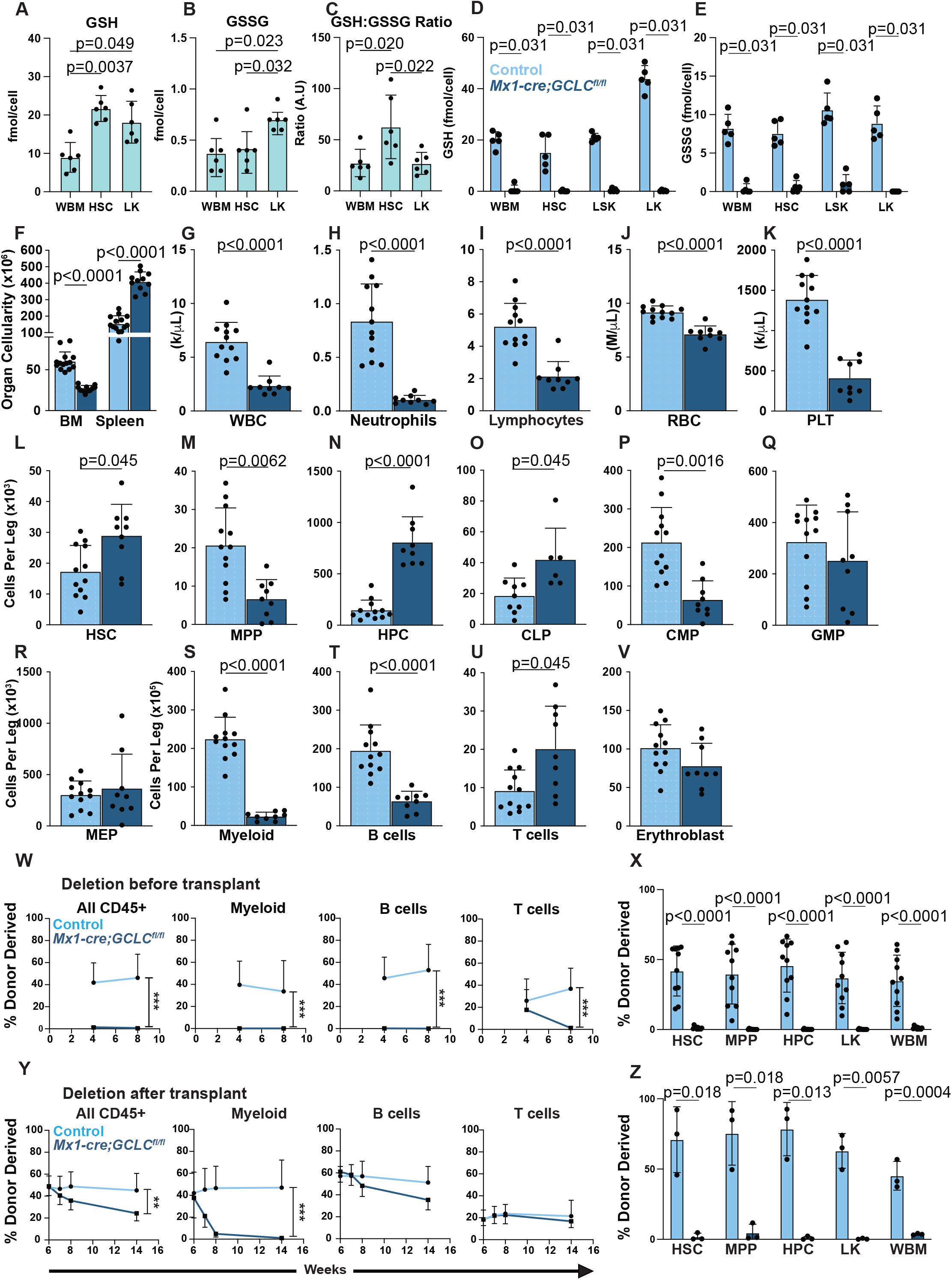
*GCLC* deficiency in hematopoietic cells profoundly depleted glutathione, severely impairing HSC function and causing pancytopenia. (A-C) Reduced glutathione (GSH) (A), oxidized glutathione (GSSG) (B), and the ratio of reduced to oxidized glutathione (C) in whole bone marrow cells, HSCs and LK myeloid progenitors in normal wild-type mice (n=6 mice in two independent experiments). (D-E) The levels of GSH (D) and GSSG (E) in control and *Mx1-cre;GCLC^fl/fl^* hematopoietic cells (n=5 mice in two independent experiments). (F) Bone marrow (one femur and one tibia) and spleen cellularity in *Mx1-cre;GCLC^fl/fl^*and littermate control mice (n=9-12 mice per genotype from three independent experiments). (G-K) White blood cell, neutrophil, lymphocyte, red blood cell, and platelet counts in *Mx1-cre;GCLC^fl/fl^* and littermate control mice (n=9-12 mice per genotype from three independent experiments). (L-V) Numbers of HSCs (L), MPPs (M), HPCs (N), CLPs (O), CMPs (P), GMPs (Q), MEPs (R), Mac1^+^CD11b^+^ Myeloid cells (S), B220^+^ B cells (T), CD3^+^ T cells (U), and CD71^+^Ter119^+^ basophilic erythroblasts (V) in the bone marrow from one tibia and one femur of *Mx1-cre;GCLC^fl/fl^* and littermate control mice (n=6-12 mice per genotype from three independent experiments). (W-X) Donor cell contributions to CD45^+^ cells, Mac1^+^CD11b^+^ myeloid cells, B220^+^ B cells, and CD3^+^ T cells in the blood (W) and HSCs, MPPs, HPCs, LK cells, and whole bone marrow cells in the bone marrow (Y) of primary recipient mice competitively transplanted with *Mx1-cre;GCLC^fl/fl^* or littermate control donor bone marrow two weeks after treatment with polyI:C to induce recombination (2 donors per genotype were transplanted into 10 recipient mice per genotype in 2 independent experiments). (Y-Z) Donor cell contributions in the blood (Y) and bone marrow (Z) of primary recipient mice competitively transplanted with *Mx1-cre;GCLC^fl/fl^*or littermate control donor bone marrow, followed by polyI:C treatment 6 weeks after transplantation to induce recombination (2 donors per genotype were transplanted into 6-10 recipient mice per genotype in 2 independent experiments). Each point represents a different mouse, except in panels W and Y. All data represent mean ± standard deviation. Statistical significance was assessed using matched samples two-way ANOVAs followed by Sidak’s multiple comparisons adjustments (**a,b,f**), a one-way ANOVA followed by Tukey’s multiple comparisons adjustment (**c**), Student’s *t*-tests, Welch’s *t*-tests, or Mann-Whitney tests followed by Holm-Sidak’s multiple comparisons adjustments (**d, e, g-k, l-v, x,** and **z**), or nonparametric analyses of longitudinal data (nparLDs) followed by Benjamini-Hochberg’s False Discovery Rate (FDR) method for multiple comparisons adjustments (**w** and **y**). All statistical tests were two-sided.

To test if glutathione is necessary for HSC function or hematopoiesis, we generated *Mx1-cre; GCLC^fl/fl^* mice and treated with four doses of polyinosinic-polycitidilic acid (poly I:C) at 6-8 weeks of age to induce recombination. Two to three weeks after the last dose of poly I:C, we isolated HSCs, Lineage^-^Sca1^+^c-kit^+^ (LSK) hematopoietic stem and progenitor cells, LK myeloid progenitors, and unfractionated bone marrow cells from *Mx1-cre; GCLC^fl/fl^* and littermate control mice and performed targeted metabolomic analysis of GSH and GSSG. All samples from *Mx1-cre; GCLC^fl/fl^* mice exhibited little or no GSH or GSSG (Figure 1D-E). GCLC was thus necessary for glutathione in hematopoietic stem and progenitor cells.

*Mx1-cre; GCLC^fl/fl^* mice were pancytopenic 2-3 weeks after poly I:C treatment, marked by significantly reduced cellularity in the bone marrow (Figure 1F) and white blood cell, neutrophil, lymphocyte, red blood cell, and platelet counts in the blood (Figure 1G-K). Spleen cellularity was significantly increased in *Mx1-cre; GCLC^fl/fl^* as compared to control mice (Figure 1F), suggesting an induction of extramedullary hematopoiesis to compensate for bone marrow and blood cell depletion. Based on surface marker phenotypes, the numbers of some hematopoietic stem and progenitor cells were depleted in the bone marrow, including multipotent progenitors (MPPs; Figure 1M), common myeloid progenitors (CMPs; Figure 1P) as well as differentiated myeloid cells (Figure 1S) and B cells (Figure 1T). Other hematopoietic stem and progenitor cells were present in normal or elevated numbers in the bone marrow based on surface marker phenotypes, including HSCs (Figure 1L), CD48^+^Lineage^-^Sca1^+^c-kit^+^ hematopoietic progenitor cells (HPCs; Figure 1N), common lymphoid progenitors (CLPs, Figure 1O), and differentiated T cells (Figure 1U).

To test HSC function in *Mx1-cre; GCLC^fl/fl^* mice, we performed a series of competitive transplantation assays. First, we treated *Mx1-cre; GCLC^fl/fl^* and littermate control mice with poly I:C, then 2-3 weeks later competitively transplanted 500,000 unfractionated bone marrow cells from these donor (CD45.2^+^) mice in competition with 500,000 recipient (CD45.1^+^) bone marrow cells into irradiated recipient mice. *Mx1-cre; GCLC^fl/fl^* bone marrow cells gave little or no donor cell reconstitution in the myeloid or B lineages, and T cell reconstitution that declined over time (Figure 1W). After not observing multilineage donor cell reconstitution in any of the recipients of *Mx1-cre; GCLC^fl/fl^*cells, we killed the mice 8 weeks after transplantation and examined donor cell reconstitution in the bone marrow. Again, we observed little or no contribution of *Mx1-cre; GCLC^fl/fl^* cells to HSCs, MPPs, HPCs, LK cells, or whole bone marrow cells in recipient bone marrow (Figure 1X). GCLC is thus necessary HSC function.

The decline in HSC function after *GCLC* deletion could have reflected a cell autonomous requirement for GCLC in HSCs or a non-cell-autonomous effect, such as exhaustion caused by HSC proliferation in response to pancytopenia. To test this, we transplanted *Mx1-cre; GCLC^fl/fl^* or control bone marrow cells, along with wild-type competitors, into irradiated recipient mice, allowed them to engraft for 6 weeks and then treated with poly I:C to induce *GCLC* deletion. Within weeks of poly I:C treatment, we observed a dramatic decline in donor cell reconstitution in recipients of *Mx1-cre; GCLC^fl/fl^*, but not control, donor bone marrow cells (Figure 1Y). All, or nearly all, donor myeloid cells were lost within 8 weeks of poly I:C treatment in recipients of *Mx1-cre; GCLC^fl/fl^* cells. Consistent with this, we observed little or no contribution of *Mx1-cre; GCLC^fl/fl^* cells to the reconstitution of HSCs, MPPs, HPCs, LK cells, or whole bone marrow cells in recipient mouse bone marrow (Figure 1Z). The absence of *GCLC*-deficient HSC function in this experiment cannot be explained by a non-cell-autonomous effect of pancytopenia because the presence of competing wild-type cells in these recipient mice prevented the development of pancytopenia after poly I:C treatment (Suppl. Figure 1B). GCLC was thus cell-autonomously required for HSC function.

### Glutathione deficiency increases reactive oxygen species in hematopoietic cells

To test if the loss of HSC function in *Mx1-cre; GCLC^fl/fl^*mice was associated with elevated ROS levels, we stained *Mx1-cre; GCLC^fl/fl^*and control bone marrow cells with Enzo total ROS dye, BODIPY C11 dye (which detects lipid peroxidation^38^), and CellRox DeepRed (which detects cytoplasmic ROS, according to the manufacturer) 2-3 weeks after poly I:C treatment. Enzo total ROS, BODIPY C11, and CellRox DeepRed staining were significantly elevated in most hematopoietic stem and progenitor cell populations we examined in *Mx1-cre; GCLC^fl/fl^* as compared to control bone marrow, including in HSCs (Figure 2A-C). To test if oxidative stress contributed to the loss of HSC function and the pancytopenia observed in *Mx1-cre; GCLC^fl/fl^* mice, we treated *Mx1-cre; GCLC^fl/fl^* and littermate control mice with poly I:C, with or without daily injections of N-acetyl-cysteine (NAC), starting when the poly I:C was administered. NAC is an antioxidant that scavenges ROS as well as increasing the availability of cysteine for glutathione synthesis, though the latter effect would not be expected in *GCLC*-deficient cells, which lack the ability to synthesize glutathione from cysteine^39,40^. NAC treatment reduced total cytoplasmic ROS levels to control levels in whole bone marrow but did not significantly rescue the depletion of bone marrow cells, the increase in spleen cells, or the decline in blood cell counts in *Mx1-cre; GCLC^fl/fl^* as compared to littermate control mice (Figure 2D-2H).

**Figure 2.**
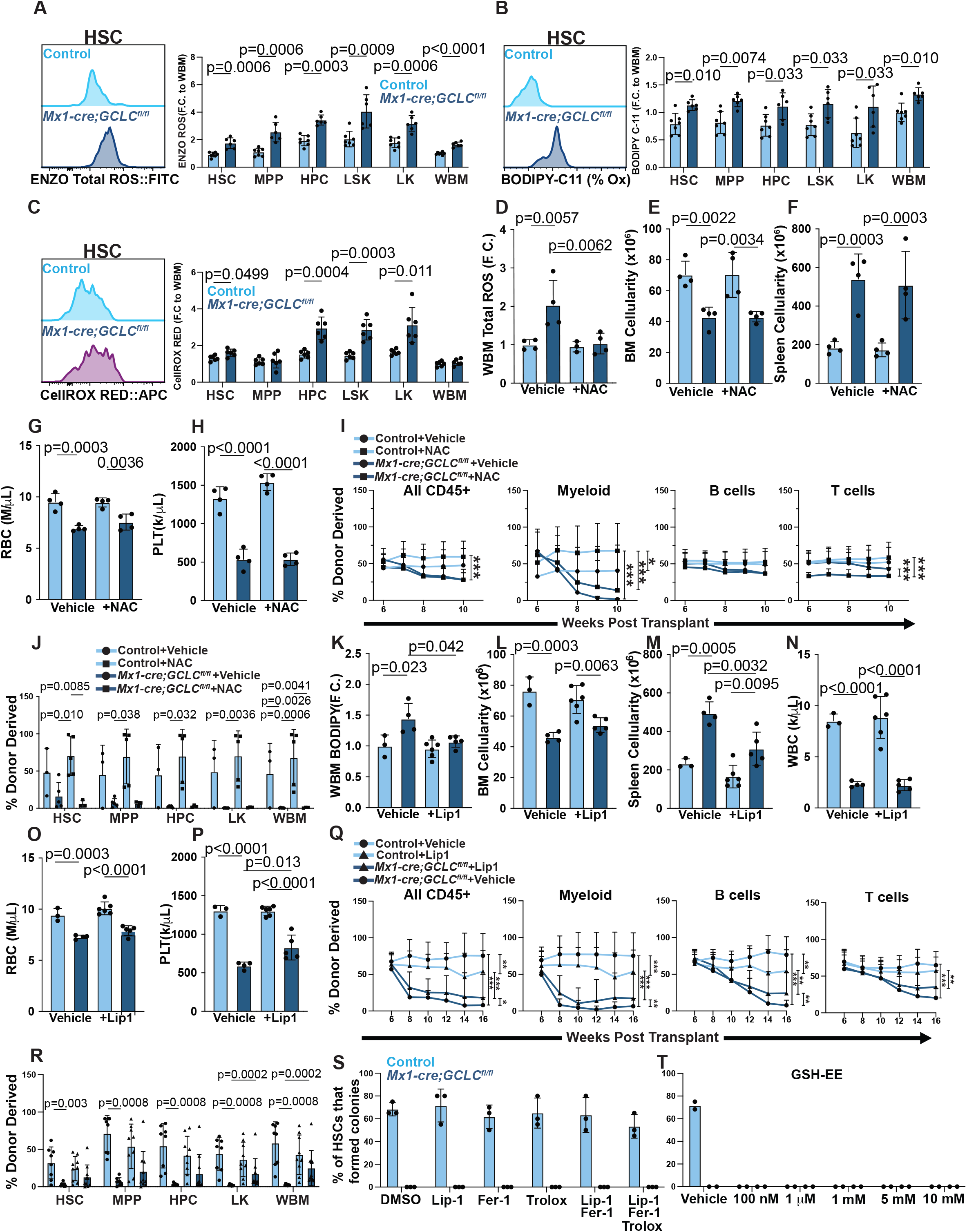
*GCLC* deficiency increased ROS levels in hematopoietic stem and progenitor cells but treatment with antioxidants and/or ferroptosis inhibitors gave little or no rescue of HSC function or pancytopenia. (A-C) Representative histograms for ENZO total ROS (A), BODIPY C-11 (B), and CellROX Deep Red (C) staining of HSCs as well as fold change in median fluorescence intensity between *Mx1-cre; GCLC^fl/fl^* versus control HSCs, MPPs, HPCs, LSKs, LKs, and whole bone marrow cells, normalized to whole bone marrow cells in each experiment (a total of 6-7 mice per genotype in three independent experiments for **A** and **B**, and 4 mice per genotype in two independent experiments for **C**). (D) Fold change in median fluorescence intensity of ENZO Total ROS staining for *Mx1-cre; GCLC^fl/fl^* versus control whole bone marrow cells from N-acetyl-cysteine or vehicle-treated mice (a total of 4 mice per treatment from two independent experiments). (E-H) Bone marrow (E) and spleen (F) cellularity as well as red blood cell (G) and platelet (H) counts from *Mx-1-cre; GCLC^fl/fl^*and littermate control mice treated with N-acetyl-cysteine or vehicle (a total of 4 mice per treatment from two independent experiments). (I-J) Donor cell contributions to the blood (I) and bone marrow (J) of mice competitively transplanted with *Mx1-cre; GCLC^fl/fl^* or control donor bone marrow cells, followed by poly I:C treatment 6 weeks after transplantation to induce recombination. The recipient mice received injections of N-acetyl-cysteine or vehicle every other day until endpoint (1 donor per genotype was transplanted into 3-5 recipient mice per treatment). (K) Fold change in median fluorescence intensity of BODIPY C-11 oxidation staining for *Mx1-cre; GCLC^fl/fl^* versus control bone marrow cells from liproxstatin-1 or vehicle-treated mice (a total of 3-6 mice per treatment from two independent experiments). (L-P) Bone marrow (L) and spleen (M) cellularity as well as white blood cell (N) red blood cell (O) and platelet (P) counts from *Mx-1-cre; GCLC^fl/fl^* and littermate control mice treated with liproxstatin-1 or vehicle (a total of 3-6 mice per treatment from two independent experiments). (Q-R) Donor cell contributions to the blood (Q) and bone marrow (R) of mice competitively transplanted with *Mx1-cre; GCLC^fl/fl^*or control donor bone marrow cells, followed by poly I:C treatment 6 weeks after transplantation to induce recombination. The recipient mice received injections of liproxstatin-1 or vehicle every day until endpoint (2 donors per genotype were transplanted into 8-10 mice per treatment in 2 independent experiments). (S) Colony formation by HSCs from *Mx1-cre; GCLC^fl/fl^* and control mice cultured in DMSO (vehicle), 1uM Liprostatin-1, 1uM Ferrostatin-1, and/or 300 uM Trolox (a total of 3 mice per treatment from 3 independent experiments). (T) The percentage of HSCs from *Mx1-cre; GCLC^fl/fl^* and control mice that formed colonies in control cultures (vehicle) or cultures supplemented by 100nM, 1uM, 1mM, 5mM, or 10 mM glutathione ethyl-ester (a total of 2 mice per treatment from 2 independent experiments). Each dot represents a different mouse, except in panels I and Q. All data represent mean ± SD. Statistical significance was assessed using Student’s *t*-tests, Welch’s *t*-tests, or Mann-Whitney tests followed by Holm-Sidak’s multiple comparisons adjustments (**a**-**c,** and **r,** and **s**), one-way ANOVAs followed by Sidak’s multiple comparisons adjustments (**d-h** and **j-p**), Kruskal-Wallis tests followed by Dunn’s multiple comparisons adjustments (**j**), or nparLDs followed by FDR multiple comparisons adjustments (**i,q**). All statistical tests were two-sided.

To test if NAC administration rescued HSC function, we competitively transplanted *Mx1-cre; GCLC^fl/fl^*and control donor cells into irradiated recipient mice, allowed them to engraft for 6 weeks, and then treated with poly I:C to induce *GCLC* recombination. We treated half of the mice in each group with NAC injections every second day for four weeks. *Mx1-cre; GCLC^fl/fl^* and control bone marrow cells gave similar levels of donor cell reconstitution until poly I:C was administered. At that point, recipients of *Mx1-cre; GCLC^fl/fl^* bone marrow cells exhibited a significant decline in donor cell reconstitution, including the loss of all, or nearly all, donor myeloid cells over a four-week period (Figure 2I). Consistent with this, we observed little contribution of *Mx1-cre; GCLC^fl/fl^*bone marrow cells to HSC, MPP, HPC, LK cell, or whole bone marrow cells in recipient mice 4 weeks after poly I:C treatment (Figure 2J). NAC treatment did not significantly affect donor cell reconstitution in the blood or the bone marrow of recipients of *Mx1-cre; GCLC^fl/fl^* or control bone marrow cells (Figure 2I and 2J). NAC treatment, therefore, did not rescue the function of *Mx1-cre; GCLC^fl/fl^*HSCs.

Since the antioxidant effects of NAC depend partly on increased glutathione synthesis^39^ and this would be blocked by *GCLC* deficiency, we also tested if liproxstatin-1 (Lip1) injection every day rescued HSC function or hematopoiesis in *Mx1-cre; GCLC^fl/fl^* mice. Liproxstatin-1 is a radical-trapping antioxidant that inhibits ferroptosis by reducing lipid peroxides^41^. Bodipy C11 staining was significantly increased in the bone marrow of *Mx1-cre; GCLC^fl/fl^* as compared to control mice and this was rescued by liproxstatin-1 treatment (Figure 2K). Nonetheless, liproxstatin-1 treatment did not rescue the pancytopenia in *Mx1-cre; GCLC^fl/fl^* mice. Irrespective of whether the mice were treated with liproxstatin-1, *Mx1-cre; GCLC^fl/fl^* mice had significantly reduced bone marrow cellularity (Figure 2L) and blood cell counts relative to control mice (Figure 2N-P). *Mx1-cre; GCLC^fl/fl^* mice had significantly increased spleen cellularity as compared to control mice, though the magnitude of the increase was reduced by liproxstatin treatment (Figure 2M). Liproxstatin-1, therefore, did not rescue the reduced bone marrow and blood cellularity in *GCLC*-deficient mice but may have attenuated the increase in extramedullary hematopoiesis.

To test if liproxstatin-1 rescued HSC function, we competitively transplanted *Mx1-cre; GCLC^fl/fl^* and control donor cells into irradiated mice, allowed them to engraft for 6 weeks, and then treated with poly I:C to delete *GCLC*. We treated half of the mice in each group with injections of liproxstatin-1 every day for ten weeks. *Mx1-cre; GCLC^fl/fl^*and control bone marrow cells gave similar levels of donor cell reconstitution until poly I:C was administered, then recipients of *Mx1-cre; GCLC^fl/fl^* cells exhibited a significant decline in donor cell reconstitution as compared to control donor cells (Figure 2Q). Consistent with this, we observed significantly lower contributions of *Mx1-cre; GCLC^fl/fl^* as compared to control bone marrow cells to HSC, MPP, HPC, LK cell, and whole bone marrow cells in recipient mice 10 weeks after poly I:C treatment (Figure 2R). Liproxstatin-1 treatment significantly increased donor cell reconstitution by *Mx1-cre; GCLC^fl/fl^* cells but not control bone marrow cells (Figure 2Q and 2R). This was evident among myeloid and B cells in the blood and LK cells and unfractionated cells in the bone marrow; however, the magnitude of the increase was small.

To further explore the effect of antioxidants and ferroptosis inhibitors on the function of *GCLC*-deficient HSCs, we sorted HSCs into cultures supplemented by liproxstatin-1, ferrostatin-1, and/or Trolox (a vitamin E analogue) and assessed the effects on colony formation. Most control HSCs formed colonies, irrespective of antioxidants, but *GCLC*-deficient HSCs consistently failed to form colonies, even in the presence of some or all of these antioxidants (Figure 2S). We also tested whether addition of 0.1, 1, 5, or 10mM of cell permeable glutathione ethyl ester to the culture medium would rescue colony formation by *GCLC*-deficient HSCs; however, neither control nor *GCLC*-deficient HSCs formed colonies in cultures supplemented with glutathione ethyl ester, precluding any conclusion (Figure 2T). These results suggest that ferroptosis or oxidative stress may have contributed to the loss of *GCLC*-deficient HSC function but that most of the defect in HSC function was not rescued by antioxidant treatment.

### GPX4 is dispensable for HSC function in vivo

GPX4 is a key cellular defense against ferroptosis: it employs glutathione to reduce lipid peroxides and to prevent the propagation of lipid oxidation within membranes^42^. To assess GPX4 function in HSCs, we analyzed *Vav-iCre; GPX4^fl/fl^* and littermate control mice at 8 to 10 weeks of age. *GPX4*-deficient and control HSCs, MPPs, HPCs, LSK cells, LK cells, and unfractionated bone marrow cells did not significantly differ in terms of ENZO total ROS staining (Suppl. Figure 2A) but all of these cell populations had significantly higher levels of Bodipy C11 staining (Suppl. Figure 2B), suggesting higher levels of lipid ROS. Consistent with a prior study^20^, *GPX4*-deficient HSCs did not form colonies in culture, unless Trolox was added to the cultures (Suppl. Figure 2C); however, colony formation was not rescued by the ferroptosis inhibitors liproxstatin-1 or ferrostatin-1 in our study or the prior study^20^. HSCs thus require GPX4 to limit lipid peroxidation in vivo and for colony formation in culture though it is not clear whether the lack of colony formation by *GPX4*-deficient HSCs is caused by ferroptosis.

To test if GPX4 is necessary for HSC function in vivo, we analyzed hematopoiesis in *Vav-iCre; GPX4^fl/fl^* and littermate control mice at 8 to 10 weeks of age. *Vav-iCre; GPX4^fl/fl^* and control mice had similar bone marrow cellularity (Suppl. Figure 2D), white blood cell (Suppl. Figure 2E), neutrophil (Suppl. Figure 2F), and platelet (Suppl. Figure 2I) counts but the *Vav-iCre; GPX4^fl/fl^* mice had significantly lower lymphocyte (Suppl. Figure 2G) and red blood cell (Suppl. Figure 2H) counts, consistent with the known requirements for GPX4 in terminal erythrocyte differentiation and T cell proliferation^21,22,43^. Perhaps due to the reduced lymphocyte and red cell counts, the *Vav-iCre; GPX4^fl/fl^* mice also had enlarged spleens relative to control mice, reflecting extramedullary hematopoiesis (Suppl. Figure 2D). Within the bone marrow, the numbers of HSCs and most restricted hematopoietic progenitors did not significantly differ between *Vav-iCre; GPX4^fl/fl^* and control mice (Suppl. Figure 2J-O); however, the numbers of MEPs (Suppl. Figure 2P) were significantly higher in *Vav-iCre; GPX4^fl/fl^* as compared to control mice, perhaps in an effort to compensate for the defect in erythrocyte differentiation.

To assess HSC function, we competitively transplanted *Vav-iCre; GPX4^fl/fl^* and control bone marrow cells into irradiated recipient mice. The *GPX4*-deficient bone marrow cells gave significantly lower levels of B and T, but not myeloid, cell reconstitution in the blood as compared to control cells (Suppl. Figure 2U). Sixteen weeks after transplantation, *GPX4*-deficient and control bone marrow cells gave rise to similar frequencies of donor HSCs, MPPs, HPCs, LK cells, and unfractionated cells in the bone marrow (Suppl. Figure 2V). Upon secondary transplantation, *GPX4*-deficient and control bone marrow cells did not significantly differ in the levels of donor cells they gave rise to in the blood or bone marrow of recipient mice (Suppl. Figure 2W and 2X). *GPX4* deficiency was thus not required for HSC function under steady-state conditions or after transplantation into irradiated mice despite reducing lipid peroxidation in HSCs. GPX4 is thus not required to limit ferroptosis in normal HSCs in vivo.

A prior study reported that *GPX4* deficiency does lead to HSC depletion in vivo if mice are fed a low vitamin E diet, though this was not confirmed with functional assays^20^. To assess this, we fed *Vav-iCre; GPX4^fl/fl^* and littermate control mice a diet with 10-fold lower vitamin E as compared to standard mouse chow (the same diet used in ref^20^). Three weeks of this diet reduced vitamin E levels in the blood to about 30% of the levels observed in mice fed standard chow (Suppl. Figure 3A). The mice could be kept on this diet long-term without exhibiting any obvious illness. Bone marrow cellularity (Suppl. Figure 3B) and red blood cell counts (Suppl. Figure 3F) were significantly reduced in *GPX4*-deficient as compared to control mice fed a low vitamin E diet, though other blood cell counts did not significantly differ (Suppl. Figure 3C-E, G). HSC frequency in the bone marrow did not significantly differ between *GPX4*-deficient and control mice on a low vitamin E diet (Suppl. Figure 3H) but the absolute number of HSCs in the bone marrow of the *GPX4*-deficient mice was significantly reduced because overall bone marrow cellularity was reduced (Suppl. Figure 3I). The numbers of restricted hematopoietic progenitors in the bone marrow did not significantly differ between *GPX4*-deficient and control mice on a low vitamin E diet (Suppl. Figure 3J-O). However, the numbers of T cells in the bone marrow were lower in *GPX4*-deficient as compared to control mice (Suppl. Figure 3P-S), consistent with the requirement for *GPX4* during T cell proliferation^43^.

To test the effect of *GPX4* deficiency on HSC function in the context of a low vitamin E diet we competitively transplanted *Vav-iCre; GPX4^fl/fl^* and control bone marrow cells from mice fed a low vitamin E diet into irradiated recipient mice that were also fed a low vitamin E diet. Consistent with the data on HSC frequency (Suppl. Figure 3H), we observed no significant differences between *GPX4*-deficient and control bone marrow cells in terms of the levels of donor cell reconstitution in the blood or bone marrow, except for T cells (Suppl. Figure 3T and 3U). Therefore, *GPX4* was not required for normal HSC function, even when donor and recipient mice were fed a low vitamin E diet, though the low vitamin E diet may have accentuated the effects of *GPX4* deficiency on red cell differentiation and T cell proliferation.

**Figure 3.**
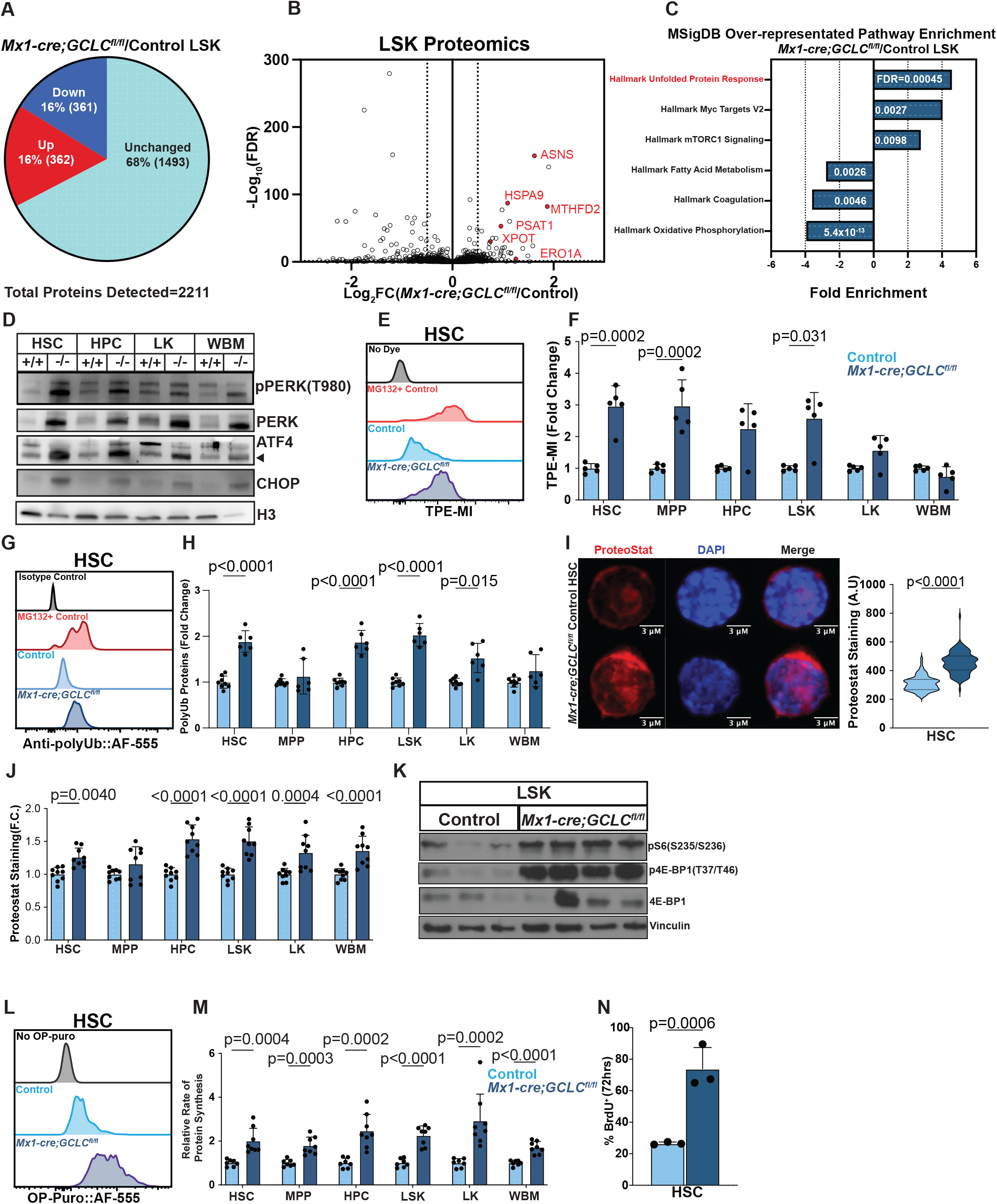
Glutathione depletion causes proteostatic stress in hematopoietic stem and progenitor cells. (A-C) We performed tandem-mass tagged proteomics on LSK cells from *Mx1-cre; GCLC^fl/fl^* or littermate control mice. We detected 2178 to 2202 proteins per sample with 16% of proteins upregulated and 16% of downregulated in the mutant cells (A; FDR<0.05; n=3 mice per genotype in one experiment). (B) Volcano plot of the proteomics data. Dotted lines are fold change cutoffs (FDR<0.05, F.C>1.5). Proteins highlighted in red appear are involved in unfolded protein responses. (C) MSigDB over-representation pathway analysis of the proteomics data. (D) Western blot of 9,000 sorted HSCs, HPCs, LK, and WBM cells per lane from *Mx1-cre; GCLC^fl/fl^* and control mice for phosphorylated PERK (T980), total PERK, ATF4, and CHOP relative to Histone3 loading control (1 mouse per genotype from one experiment; results are representative of 3 independent experiments). (E-F) Representative histograms for tetraphenylethene malemide fluorescence (E) in HSCs as well as fold change in median fluorescence intensity between *Mx1-cre; GCLC^fl/fl^* versus control HSCs, MPPs, HPCs, LSKs, LKs, and whole bone marrow cells, normalized to whole bone marrow cells in each experiment (a total of 5 mice per genotype in two independent experiments). (G-H) Representative histograms for polyubiquitinated protein staining (G) in HSCs as well as fold change in median fluorescence intensity between *Mx1-cre; GCLC^fl/fl^* versus control HSCs, MPPs, HPCs, LSKs, LKs, and whole bone marrow cells (a total of 6-8 mice per genotype in two independent experiments). (I) Representative confocal microscopy images of Proteostat staining in HSCs from *Mx1-cre; GCLC^fl/fl^* and control mice along with quantitation of staining intensity (HSCs are from 2 mice per genotype in two independent experiments). (J) Fold change in median fluorescence intensity of Proteostat staining in HSCs, MPPs, HPCs, LSKs, LKs, and whole bone marrow cells from *Mx1-cre; GCLC^fl/fl^* versus control mice (9 mice per genotype from three independent experiments). (K) Western blot of 100,000 sorted LSK cells per lane from *Mx1-cre; GCLC^fl/fl^* and control mice for phosphorylated S6 (S235/S236), phosphorylated 4E-BP1 (T37/T46), and total 4E-BP1 relative to vinculin loading control (one mice per lane from one experiment). (L-M) Representative histograms of O-propargyl-puromycin incorporation (L) in HSCs as well as fold change in median fluorescence intensity between *Mx1-cre; GCLC^fl/fl^*versus control HSCs, MPPs, HPCs, LSKs, LKs, and whole bone marrow cells (a total of 7-8 mice per genotype in two independent experiments). (N) The percentage of HSCs that incorporated a 72-hour pulse of BrdU in *Mx1-cre;GCLC^fl/fl^* and littermate control mice (3 mice per genotype). Each point represents a different mouse, except in panels B and I. All data represent mean ± standard deviation. Statistical significance was assessed using Student’s *t*-tests, Welch’s *t*-tests, or Mann-Whitney tests followed by Holm-Sidak’s multiple comparisons adjustments (**f**, **h**, **j**, and **n**), a Student’s *t*-test (**i**), or a matched samples two-way ANOVA followed by Sidak’s multiple comparisons adjustment (**m**). All statistical tests were two-sided.

### *GCLC*-deficient HSCs exhibit proteostatic stress

Since HSCs and most restricted hematopoietic progenitors did not require *GPX4* and we could not substantially rescue HSC function in *Mx1-cre; GCLC^fl/fl^* mice by treating with antioxidants and/or ferroptosis inhibitors, we wondered if *GCLC*-deficient HSCs exhibited other defects. To test this, we performed proteomic analysis in LSK cells from *Mx1-cre; GCLC^fl/fl^* and littermate control mice two weeks after treating with poly I:C. We detected 2178 to 2202 proteins in each sample: 16% of proteins were significantly (FDR<0.05) less abundant and 16% were significantly more abundant in LSK cells from *Mx1-cre; GCLC^fl/fl^*as compared to control mice (Figure 3A). Several of the proteins that were most significantly upregulated in *GCLC*-deficient as compared to control LSK cells were ATF4 transcriptional targets involved in the integrated stress response, including Asparagine synthetase (Asns^44^), HSPA9 (a mitochondrial Hsp70 chaperone^45^), mitochondrial methylenetetrahydrofolate dehydrogenase 2 (MTHFD2^46^), phosphoserine aminotransferase 1 (PSAT1^47^), exportin-t (XPOT, which exports tRNAS to the cytoplasm, increasing protein synthesis capacity in response to stress^48^), and ER oxidoreductin 1 alpha (ERO1A^49^) (Figure 3B). Using the genes that encode the proteins that significantly differed between *GCLC*-deficient and control LSK cells (FDR<0.05, FC>1.5), we performed MSigDB over-representation analysis. The most significantly enriched gene set in *GCLC*-deficient as compared to control LSK cells related to unfolded protein responses (Figure 3C). Other enriched gene sets related to Myc targets and mTORC1 signaling while gene sets related to oxidative phosphorylation, fatty acid metabolism, and coagulation were depleted (Figure 3C).

To further test whether *GCLC* deficiency caused proteostatic stress in hematopoietic stem and progenitor cells, we evaluated the levels of proteins involved in unfolded protein responses by western blot. *GCLC*-deficient HSCs, HPCs, LK cells, and whole bone marrow cells consistently exhibited higher levels of phosphorylated PERK (T980), total PERK, ATF4, and CHOP as compared to control cells (Figure 3D). By flow cytometry, *GCLC*-deficient HSCs, MPPs, and LSK cells, but not whole bone marrow cells, exhibited significantly higher levels of TPE-MI (tetraphenylethene maleimide) staining as compared to control cells (Figure 3E and 3F). TPE-MI fluoresces after reacting with free cysteine thiols, such that increased fluorescence is associated with increased unfolded protein and proteostatic stress^50^. We also observed significantly increased staining for polyubiquitinated proteins in *GCLC*-deficient as compared to control HSCs, HPCs, LSK cells, and LK cells, but not whole bone marrow cells (Figure 3G and 3H). Finally, ProteoStat staining, which reveals misfolded and aggregated proteins^51^, was significantly increased in *GCLC*-deficient as compared to control HSCs, HPCs, LSK cells, LK cells, and whole bone marrow cells (Figure 3I and 3J). Hematopoietic stem and progenitor cells thus exhibited many signs of proteostatic stress after *GCLC* deletion.

Proteostatic stress initially attenuates translation, but subsequently selectively increases the translation of subsets of mRNAs that promote adaptation to proteostatic stress^52–54^, including Myc and mTORC1 targets^55^. Consistent with this, western blot analysis showed increased mTORC1 signaling in *GCLC*-deficient as compared to control LSK cells marked by higher levels of phosphorylated S6 (S235/S236), phosphorylated 4E-BP1 (S65) and total 4E-BP1 (Figure 3K). We also assessed rates of protein synthesis by administering a 1 hour pulse of OP-Puro to *Mx1-cre; GCLC^fl/fl^*and control mice. OP-Puro incorporates into nascent peptides, such that staining correlates with the rate of protein synthesis^28,56^. We observed significantly increased OP-Puro incorporation into *GCLC*-deficient as compared to control HSCs, MPPs, HPCs, LSK cells, LK cells, and whole bone marrow cells (Figure 3L and 3M). *GCLC*-deficient hematopoietic stem and progenitor cells thus exhibited increased mTORC1 staining and protein synthesis as compared to control cells (Figure 3C). Consistent with the increased mTORC1 staining, Myc target expression, and protein synthesis, we also observed increased cell division by *GCLC*-deficient as compared to control HSCs (Figure 3N), potentially explaining why the number of HSCs in the bone marrow based on surface marker phenotype increased in the first few weeks after *GCLC* deletion (Figure 1L).

In contrast to the proteostatic stress observed in hematopoietic stem and progenitor cells after *GCLC* deletion, we did not observe signs of proteostatic stress in *GPX4*-deficient HSCs. HSCs, MPPs, HPCs, LSK cells, LK cells, and whole bone marrow cells from *Vav-icre;GPX4^fl/fl^* and littermate control mice did not significantly differ in terms of TPE-MI (unfolded protein) staining, polyubiquitinated protein levels, proteostat staining, or levels of proteins involved in unfolded protein responses (Suppl. Figure 4A-D). Therefore, the increase in proteostatic stress in *GCLC*-deficient as compared to control hematopoietic stem and progenitor cells seemed to arise independently of the ability of cells to use glutathione to block lipid peroxidation.

### *p53* deletion prolonged the function of *GCLC*-deficient HSCs after transplantation

Sustained proteostatic stress can induce p53-mediated apoptosis^57^. Consistent with this, we observed substantially increased levels of p53 in LSK cells from *Mx1-cre; GCLC^fl/fl^* as compared to control mice (Figure 4A). We also observed significantly increased levels of Annexin V staining among propidium iodide negative HPCs and LSK cells, as well as a trend toward increased Annexin V staining in MPPs, LK cells, and whole bone marrow cells from *Mx1-cre; GCLC^fl/fl^* as compared to control mice (Figure 4B). Increased Annexin V staining in cells with membrane integrity (propidium iodide negative) is a marker of early apoptosis^58^, though Annexin V staining increases after the loss of membrane integrity in cells that undergo multiple types of cell death^59^. *p53* deficiency significantly reduced the percentages of MPPs, HPCs, LSK cells, LK cells, and whole bone marrow cells that were Annexin V+ and propidium iodide negative in *Mx1-cre; GCLC^fl/fl^* mice (Figure 4B). These data suggested that *GCLC* deficiency caused some stem and progenitor cells to undergo p53-mediated cell death, perhaps by apoptosis.

**Figure 4.**
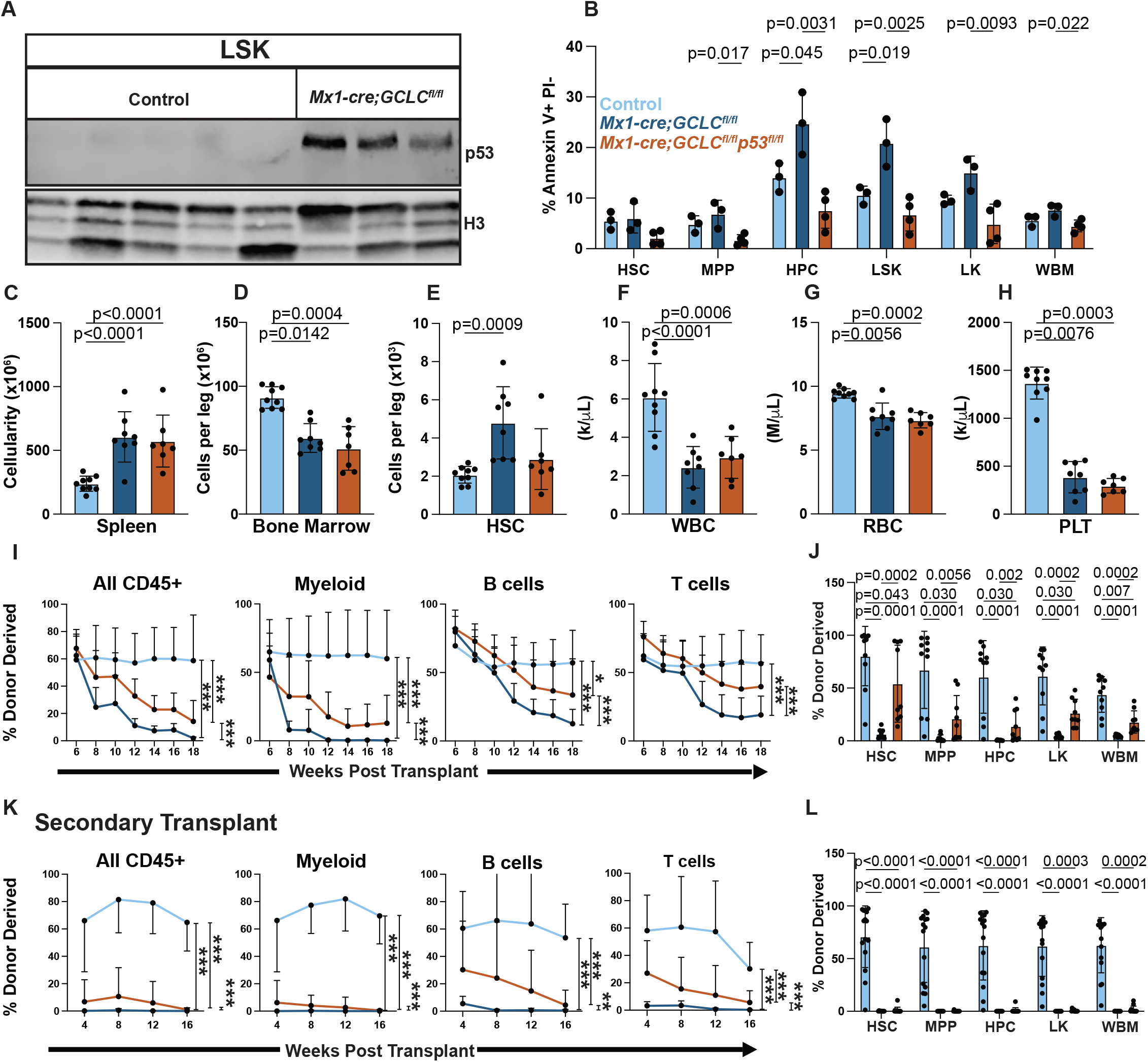
The loss of HSC function after *GCLC* deletion was marked by increased p53 levels and Annexin V staining. (A) Western blot of total p53 levels in 100,000 LSK cells per lane from *Mx1-cre; GCLC^fl/fl^* and controls mice relative to Histone3 loading control (one mouse per lane; results are representative of 2 independent experiments). (B) Percentage of HSCs, MPPs, HPCs, LSKs, LKs, and whole bone marrow cells from *Mx1-cre; GCLC^fl/fl^* mice, *Mx1-cre; GCLC^fl/fl^; p53^fl/fl^* mice, and littermate controls that stained positively for Annexin V and negatively for propidium iodide (PI) (3-4 mice per genotype from 1 experiment). (C-H) Spleen (C) and bone marrow (D) cellularity as well as HSC number in the bone marrow (E), and white blood cell (F), red blood cell (G), and platelet (H) counts in the blood of *Mx1-cre; GCLC^fl/fl^* mice, *Mx1-cre; GCLC^fl/fl^; p53^fl/fl^* mice, and littermate controls (a total of 7-9 mice per genotype from three independent experiments). (I-J) Donor cell contributions to the blood (I) and bone marrow (J) of mice competitively transplanted with *Mx1-cre; GCLC^fl/fl^*, *Mx1-cre; GCLC^fl/fl^; p53^fl/fl^*, or control donor bone marrow, followed by poly I:C treatment 6 weeks after transplantation to induce recombination (2 donors per genotype transplanted into 9-10 recipient mice per genotype in 2 independent experiments). (K-L) Donor cell contributions to the blood (K) and bone marrow (L) of secondary recipient mice transplanted with bone marrow cells from the primary recipient mice in Figure 4I (2-3 primary recipient mice per genotype transplanted into 10-15 secondary recipient mice per genotype). Each dot represents a different mouse except in panels I and K. All data represent mean ± SD. Statistical significance was assessed using one-way ANOVAs followed by Tukey’s multiple comparisons adjustments (**b** and **f**), a Welch’s one-way ANOVA followed by Dunnett’s T3 method for multiple comparisons adjustments (**g**), Kruskal-Wallis tests followed by Dunn’s multiple comparisons adjustment (**h** and **l**), a matched samples two-way ANOVA followed by Sidak’s multiple comparisons adjustment (**c** and **d**), or nparLDs followed by FDR multiple comparisons adjustments (**i** and **k**) or Mann-Whitney tests followed by Holm-Sidak’s multiple comparisons adjustments (**e** and **j**). All statistical tests were two-sided.

To test if *p53* deficiency rescued aspects of HSC function or hematopoiesis after *GCLC* deletion, we assessed the frequencies of hematopoietic stem and progenitor cells in *Mx1-cre; GCLC^fl/fl^* and *Mx1-cre; GCLC^fl/fl^; p53^fl/fl^* as compared to control mice. Consistent with the data in Figure 1F, *GCLC* deficiency increased spleen cellularity, reduced bone marrow cellularity, and increased HSC number in the bone marrow relative to control mice (Figures 4C-4E). *p53* deficiency did not significantly alter these phenotypes (Figures 4C-4E). *GCLC* deficiency also significantly reduced white blood cell, red blood cell, and platelet counts relative to control mice, and *p53* deficiency did not significantly alter these phenotypes either (Figures 4F-4H).

To test if *p53* deficiency cell-autonomously rescued HSC function, we competitively transplanted whole bone marrow cells from *Mx1-cre; GCLC^fl/fl^* mice, *Mx1-cre; GCLC^fl/fl^; p53^fl/fl^*mice, and control mice into irradiated recipients and then induced recombination 6 weeks after transplantation by treating with poly I:C. At the time of poly I:C treatment, donor cells of all genotypes gave similar levels of donor cell reconstitution (see 6 weeks after transplantation in Figure 4I). However, 12 weeks after poly I:C treatment, *Mx1-cre; GCLC^fl/fl^* bone marrow cells gave significantly lower levels of donor cell reconstitution in all lineages as compared to control bone marrow cells and none of the recipients of *Mx1-cre; GCLC^fl/fl^*cells exhibited long-term multilineage reconstitution by donor cells (Figure 4I). *Mx1-cre; GCLC^fl/fl^; p53^fl/fl^* bone marrow cells gave significantly higher levels of donor cell reconstitution in all lineages as compared to *Mx1-cre; GCLC^fl/fl^* bone marrow cells, and 6 out of 10 recipients of double mutant cells exhibited multilineage reconstitution at 12 weeks after poly I:C treatment (Figure 4I). Consistent with this, recipients of *Mx1-cre; GCLC^fl/fl^* donor cells had few donor-derived HSCs, MPPs, HPCs, LK cells, or unfractionated cells in the bone marrow in contrast to high levels of donor chimerism in recipients of control cells (Figure 4J). Recipients of *Mx1-cre; GCLC^fl/fl^; p53^fl/fl^* cells had significantly higher donor cell contributions to HSCs, MPPs, HPCs, LK cells, and whole bone marrow cells as compared to recipients of *Mx1-cre; GCLC^fl/fl^* cells, but significantly lower contributions than recipients of control cells (Figure 4J). *p53* deficiency thus partially rescued the reconstituting potential of *GCLC*-deficient HSCs. This effect must have occurred at least partly at the level of HSCs because *p53* deficiency partially rescued the ability of donor bone marrow cells to reconstitute the HSC pool in recipient mice (Figure 4J).

To further test this, we performed secondary transplants of whole bone marrow cells from the recipients shown in Figure 4I. Sixteen weeks after secondary transplantation, all recipients of control donor cells were long-term multilineage reconstituted by donor cells, with high levels of donor chimerism in the blood (Figure 4K) and in the bone marrow (Figure 4L). Conversely, recipients of *Mx1-cre; GCLC^fl/fl^* donor cells exhibited little or no donor cell reconstitution, with no mice exhibiting long-term multilineage reconstituted by donor cells in the blood (Figure 4K) or in the bone marrow (Figure 4L). Recipients of *Mx1-cre; GCLC^fl/fl^; p53^fl/fl^* donor cells gave intermediate levels of donor cell reconstitution, significantly lower than in recipients of control donor cells but higher than in recipients of *Mx1-cre; GCLC^fl/fl^* donor cells in the blood (Figure 4K). Three of 15 recipients of *Mx1-cre; GCLC^fl/fl^; p53^fl/fl^* donor cells exhibited long-term multilineage reconstitution by donor cells in the blood and in the bone marrow. These mice exhibited low, but above background, donor cell contributions to multiple cell populations in the bone marrow (Figure 4L). *p53* deficiency thus prolonged the reconstituting potential of *GCLC*-deficient HSCs after competitive transplantation, enabling reconstitution of secondary recipients that was not otherwise observed from *GCLC*-deficient HSCs.

## Discussion

Our results demonstrate that glutathione is broadly required for the function of HSCs and most restricted hematopoietic progenitors, but not for all hematopoietic cells. *GCLC*-deficiency eliminated nearly all detectable glutathione from hematopoietic cells (Figure 1D and 1E). This led to a loss of HSC function, severe defects in hematopoiesis, and pancytopenia (Figure 1F-Z). Glutathione is most commonly thought of as a redox buffer that prevents oxidative stress, including as a co-factor employed by GPX4 to prevent lipid peroxidation and ferroptosis^42^. Although we observed increased levels of reactive oxygen species in *GCLC*-deficient hematopoietic cells, including lipid peroxide levels (Figure 2A-C), we observed little or no rescue of HSC colony formation (Figure 2S), HSC reconstituting potential (Figure 2I, J, Q, R), or hematopoiesis (Figure 2D-H, K-P) by treating with antioxidants, ferroptosis inhibitors, or both. This suggested that glutathione does regulate redox balance in hematopoietic stem and progenitor cells, but that GCLC is not required to prevent ferroptosis in normal HSCs in vivo.

The depletion of hematopoietic stem and progenitor cells after *GCLC* deletion was associated with increased p53 levels (Figure 4A) and increased Annexin V staining (Figure 4B), suggesting that these cells may have undergone apoptosis. Consistent with this possibility, glutathione depletion^60,61^ and proteotoxic stress^37^ are known to cause apoptosis in some cells. Nonetheless, while Annexin V binding to exposed phosphatidylserine on the cell surface can be an indicator of apoptosis^62^, phosphatidylserine can also be exposed to the outer membrane during other forms of cell death^63^, particularly after the loss of membrane integrity^59^.

*GPX4*-deficiency also increased lipid peroxide levels in HSCs (Suppl. Figure 2B) but had little effect on hematopoiesis in vivo, reducing the numbers of differentiated T cells and erythrocytes, consistent with prior studies^21,22,43^. *GPX4*-deficiency did not significantly affect the number of HSCs in the bone marrow (Suppl. Figure 2J) or reconstituting potential upon serial transplantation in irradiated mice (Suppl. Figure 2U-X). In contrast to *GCLC* deficiency, colony-formation by *GPX4*-deficient HSCs was substantially rescued by treating with the antioxidant Trolox. These results suggest that HSCs require GPX4 in culture, but that GPX4 function is dispensable in normal HSCs in vivo.

We found that *GCLC*-deficient HSCs and restricted hematopoietic progenitors exhibited proteostatic stress marked by an unfolded protein response (Figure 3B-D) with elevated levels of unfolded/misfolded proteins (Figure 3E and 3F), poly-ubiquitinated proteins (Figure 3G and 3H), and protein aggregates (Figure 3I and 3J). We did not detect any of these signs of proteostatic stress in *GPX4*-deficient hematopoietic stem or progenitor cells (Suppl. Figure 4). This suggests that the most critical function of glutathione in hematopoietic stem and progenitor cells is to maintain proteostasis.

HSCs appear to be more dependent upon glutathione for proteostasis than differentiated hematopoietic cells. HSCs had almost twice as much reduced glutathione as compared to whole bone marrow cells (Figure 1B). Whole bone marrow cells also did not exhibit significantly increased levels of unfolded/misfolded proteins (Figure 3F) or poly-ubiquitinated proteins (Figure 3H), in contrast to HSCs. Whole bone marrow cells did exhibit an unfolded protein response, though with somewhat lower levels of PERK and ATF4 as compared to HSCs (Figure 3D) and an increase in protein aggregates (Figure 3J). Thus, glutathione is required for proteostasis in some differentiated hematopoietic cells but HSCs exhibit more severe proteostatic stress. This may reflect alternative proteostasis mechanisms in differentiated as compared to undifferentiated cells^64^.

Glutathione promotes proteostasis partly by enabling oxidative protein folding in the endoplasmic reticulum^65^. An imbalance in the ratio of GSH to GSSG in the endoplasmic reticulum impairs protein quality control and leads to an accumulation of misfolded proteins^66^. Nonetheless, glutathione is not necessary for the folding of all proteins^23^ and not all cells require glutathione to avoid proteostatic stress^27^. It is hard to test whether the proteostatic stress we observed in HSCs is attributable to the loss of glutathione itself versus altered redox balance in the endoplasmic reticulum; however, the observation that treatment with antioxidants and/or ferroptosis inhibitors gave little or no rescue of HSC function, depending on the assay, suggests that the proteostatic stress was not caused primarily by a change in redox balance.

HSCs tend to be depleted within days in culture^67,68^. As compared to HSCs in vivo, HSC in culture are exposed to unphysiological levels of oxygen and other nutrients, exhibiting increased mitochondrial function, cell division, and protein synthesis^33,69,70^. Inhibition of protein synthesis^33^, proteostatic stress^71^, ferroptosis^18^, and cell division^72^ helps to preserve HSC function in culture but are not sufficient to enable long-term self-renewal^67^. At least some of the factors that limit HSC maintenance in culture are thus different from the factors that limit HSC function in vivo, raising the possibility that ferroptosis is more limiting for HSC maintenance in culture than in normal HSCs in vivo. Nonetheless, ferroptosis may become limiting for HSC maintenance in vivo in certain genetic backgrounds that alter HSC function^19^.

### Limitations of Study

- Although we did not observe defects in HSC function in vivo after *GPX4* deletion, either in the context of steady-state hematopoiesis or upon transplantation into irradiated mice, other stresses could render HSCs dependent upon GPX4 in vivo.
- It remains possible that ferroptosis regulates HSCs in vivo and that enzymes other than GPX4 compensate for the loss of GPX4 by suppressing lipid oxidation.

### Experimental Model and Subject Details

We used 8-12 week old male or female *GCLC^fl^* (*Gclc^tm1a(EUCOMM)Wtsi^*; ref^73^), *Mx1-cre*^74^, *Vav1-iCre*^75^, *Gpx4^fl^* (ref^76^), and *p53^fl^* (ref^77^) mice. To induce Cre expression in Mx1-Cre mice, both Mx1-Cre and control mice were administered an intraperitoneal injection of 20 μg of polyinosinic:polycytidylic acid (poly I:C) dissolved in PBS every other day for 7 days (4 injections total) starting at 6-7 weeks of age. C57BL/Ka-Thy-1.1/Thy-1.2 (CD45.1/CD45.2) mice were used as recipients and C57BL/Ka-Thy-1.2 (CD45.1) mice were used as competitor bone marrow cells in transplantation experiments. The mutant mice listed above were maintained on a C57BL/Ka-Thy-1.1 (CD45.2) background. In experiments that involved drug treatments, N-acetyl-cysteine was administered subcutaneously every day or every second day (6mg/kg/day), depending on the experiment, and Liproxstatin-1 was administered intraperitoneally every day (10 mg/kg/day). All mice were housed in specific-pathogen-free animal care facilities at UT Southwestern Medical Center (UTSW) and fed ad libitum (4% of calories from fat; Teklad Global 16% Protein Rodent Diet; Envigo #2016). Mice were randomly assigned to treatment groups. All procedures were approved by the UTSW Institutional Animal Care and Use Committee.

## Method Details

### Flow cytometric analysis and sorting of hematopoietic cells

Bone marrow was flushed from two tibias and two femurs using staining medium (HBSS with 2% heat inactivated bovine serum) and dissociated into a single cell suspension by gently triturating with a 23-gauge needle. Spleens were mechanically dissociated by crushing and triturating. Cells were resuspended in staining medium and filtered through a 40 μm cell strainer. Cells were counted and stained with antibodies at 4°C for 30 minutes (except for anti-CD34 antibody staining, which was for 90 minutes). For staining of HSCs and progenitors, cells were stained with fluorophore-conjugated antibodies against lineage markers (CD2, CD3, CD5, CD8a, Gr1, Ter119 and B220) as well as c-kit, Sca1, CD150, and CD48, CD34, CD16/32, CD127, CD134. For analysis of differentiated cells, cells were stained with fluorophore-conjugated antibodies against Mac-1, Gr1, B220, CD43, Ter119, and CD71. The markers used to identify each cell population are in Table S1 and antibodies are listed in the key resources table. Cells were analyzed using a FACS Lyric (BD Biosciences) flow cytometer. Dead cells were identified and gated out of all analyses by including propidium iodide in the staining medium used to resuspend cells for flow cytometry. Flow cytometry data were analyzed using Flowjo (BD Biosciences). To sort HSCs, tibias, femurs, pelvises, and spines were crushed using a mortar and pestle. The bone marrow cells were then gently triturated and filtered, as above, to obtain a single cell suspension. c-kit^+^ cells were enriched using anti-c-kit paramagnetic microbeads (Miltenyi Biotec). Cells were then stained with fluorophore-conjugated antibodies against lineage markers (CD2, CD3, CD5, CD8a, B220, Ter119, and Gr1), Sca1, c-kit, CD150 and CD48. Cells were isolated by two successive rounds of sorting to ensure purity using a FACSAria II or a FACSAria Fusion flow cytometer. For blood analysis, 70-100 μl of blood was collected from the retro-orbital sinus or from the heart and mixed with 5μl of 500mM EDTA to prevent clotting. Blood cell counts were determined using a Heska Element HT5 (HESKA).

### Glutathione measurement in hematopoietic cells

We used a previously described method to perform targeted metabolomic analysis on flow cytometrically isolated cells^78^. Briefly, we crushed long bones and vertebrae on ice and enriched c-kit^+^ cells using paramagnetic beads in the cold room. Cells were sorted on a FACS Aria II sorter using a 70 µm nozzle, with 0.5x PBS sheath fluid. Five thousand hematopoietic cells were sorted directly into a solution of 80% methanol containing 0.1% formic acid as reported previously^17,79,80^. Samples were vortexed, dried and resuspended in water containing 0.1% formic acid and 50 nM ^13^C_2_,^15^N_1_-glutathione (GSH) plus 50 nM ^13^C_4_,^15^N_2_-oxidized glutathione (GSSG) (Toronto Research Chemicals). Chromatographic separation of GSH and GSSG was achieved using an HSS T3 C18 column (2.1mm x 150mm) (Waters, Milford, MA) and a binary solvent system of 0.1% formic acid in water and 0.1% formic acid in acetonitrile. GSH and GSSG were analyzed using a targeted SIM acquisition method on a Q Exactive HF-X mass spectrometer (Thermo Scientific, Bremen, GE).

GSH and GSSG were quantified using isotope dilution mass spectrometry^80,81^. Briefly, a 7-point calibration curve was constructed for GSH and GSSG containing the following concentrations of unlabelled standards: 5nM, 10nM, 20nM, 50nM, 100nM, 200nM, and 500nM. All calibration points contained 50 nM ^13^C_2_,^15^N_1_-GSH and 50 nM ^13^C_4_,^15^N_2_-GSSG. For each point in the curve, the peak area ratio of unlabeled standard to isotopically labeled internal standard was calculated and plotted against known concentrations to create a regression curve. For each experimental sample, 50nM of labelled internal standard was added to each unknown sample to determine the peak area ratio of unlabeled metabolite to internal labelled standard. Applying the linear regression model, these values were used to calculate the molar concentrations of GSH and GSSG in each sample. These molar concentrations were used to calculate the final ratios of GSH and GSSG in biological samples.

### Bone marrow reconstitution assays

Recipient mice (CD45.1/CD45.2 or CD45.1) were irradiated using an XRAD 320 X-ray irradiator (Precision X-Ray Inc.) with two doses of 540 rad at least 3 hours apart. For competitive transplantation, 500,000 unfractionated bone marrow cells from donor (CD45.2) and competitor mice (CD45.1 or CD45.1/CD45.2) were mixed and intravenously injected into recipients (CD45.1 or CD45.1/CD45.2) through the retro-orbital venous sinus. For secondary transplants, 3-5 million unfractionated bone marrow cells were injected intravenously through the retro-orbital venous sinus into 5 secondary recipient mice per donor. Every 2-4 weeks until 8-16 weeks after transplantation, 50-100 μl of blood was collected from the retro-orbital venous sinus in 500 μl of HBSS containing 10 mM EDTA (to prevent clotting). Cells were subjected to ammonium-chloride potassium chloride red cell lysis and then stained with antibodies against CD45.1, CD45.2, Mac-1, Gr1, B220, and CD3 to evaluate the levels of donor cell engraftment in the myeloid, B, and T cell lineages. Levels of donor cell engraftment were evaluated in the bone marrow of recipient mice at least 8 weeks after transplantation. Cells were resuspended in staining medium, filtered through a 40 μm cell strainer, stained with antibodies against lineage markers (CD2, CD3, CD5, CD8a, Gr1, Ter119 and B220), c-kit, Sca1, CD150, and CD48, CD34, and CD16/32 as described above, and analyzed using a FACS Lyric cytometer.

### Protein extraction and western blot analysis

Equal numbers of bone marrow cells were double sorted to ensure purity and washed twice in PBS (Cytiva, SH30256.01) then lysed on ice in Triton lysis buffer (1% Triton X-100, 40 mM HEPES, pH 7.4, 120 mM NaCl, 10 mM sodium pyrophosphate, 10 mM glycerol 2-phosphate, 50 mM NaF, 1 mM EDTA, 1 μM Microcystin-LR (Enzo life sciences, ALX-350-012-C500) and protease inhibitor cocktail (Sigma, P8340)) as previously described^82^. 4X SDS loading buffer (BioRad) was added to each sample, boiled at 80°C for 10 minutes, centrifuged for 10 minutes at 16,000 x g, and the supernatants were loaded on 4-20% Mini-PROTEAN TGX gels (Biorad, 4561094). The gels were then transferred onto 0.2 μm PVDF membranes (BioRad, 1704156) by semi-dry transfer using the BioRad TransBlot Turbo transfer apparatus. Membranes were blocked in either 5% nonfat dry milk in TBST, 5% BSA in TBST (when using phosphorylated antibodies) or Intercept blocking buffer-TBS (when detecting using Licor near-IR imaging platform) (Licor Biosciences). All primary antibodies were diluted at 1:500-1000 in blocking buffer and incubated overnight at 4°C. Secondary antibodies were diluted 1:10,000-20,000 and incubated at room temperature for 1 hour. Signals were detected using either Pierce ECL western blotting chemiluminescence (Thermo Scientific, 32016, 34094) or Licor IRdye 680 or 800 secondary antibodies using a Licor Odyssey CLX.

### Protein synthesis rate

OP-Puro (50 mg/kg; pH 6.4–6.6 in PBS) was injected intraperitoneally. One hour later, mice were euthanized. Bone marrow was collected, and 1×10^7^ cells were stained with combinations of antibodies against cell surface markers as described above. Cells were fixed in 0.5 ml of 1% paraformaldehyde (Affymetrix) in PBS for 15 min on ice. After washing in PBS, cells were permeabilized in 200 μl PBS supplemented with 3% fetal bovine serum (Sigma) and 0.1% saponin (Sigma) for 5 min at room temperature (20–25°C). The azide-alkyne cycloaddition was performed using the Click-iT Cell Reaction Buffer Kit (Life Technologies) and azide conjugated to Alexa Fluor 555 (Life Technologies) at 5 μM final concentration. After the 30 min reaction, the cells were washed twice in PBS supplemented with 3% fetal bovine serum and 0.1% saponin, then resuspended in PBS and analyzed by flow cytometry.

### Reactive oxygen species

1×10^7^ bone marrow cells were stained with antibodies against cell surface markers as described above. After antibody staining, the cells were washed twice in HBSS without serum, then stained with 100 µM verapamil in HBSS plus dyes that detect different ROS for 30 minutes at 37°C. ENZO total ROS was used at 1:1000, BODIPY C-11 was used at 1 uM. Samples were washed twice with HBSS containing 100 uM verapamil, resuspended in HBSS supplemented with propidium iodide and 100 uM verapamil, then analyzed using the FACS Lyric cytometer. Percent oxidized lipid was calculated as the median fluorescence intensity (MFI) of FITC (oxidized lipid) divided by the sum of FITC and PE (non-oxidized lipid) MFIs.

### BrdU incorporation

To assess the rate of HSC cell division, mice were administered a 72-hour pulse of BrdU by intraperitoneally injecting with 0.1 mg/g body mass of BrdU dissolved in PBS and then maintained on 1 mg/ml BrdU in the drinking water for 72 hours. Bone marrow cells from the spine and legs were stained with antibodies against HSC markers and c-kit^+^ cells were enriched using paramagnetic beads as described above. The cells were fixed and stained with anti-BrdU antibody using the BD APC BrdU Flow Kit following the manufacturer’s instructions (Becton Dickson). The percentage of BrdU+ HSCs was determined using a FACS Lyric flow cytometer.

### Unfolded, misfolded and aggregated proteins

1×10^7^ bone marrow cells were stained with antibodies against cell surface markers as described above before being stained with either of three reagents that detect unfolded or misfolded proteins. Some cells were incubated with 50 uM TPE-MI (tetraphenylethene maleimide) in 200 ul of serum-free HBSS for 45 minutes at 37°C as one measure of unfolded proteins. Other cells were fixed in 1% paraformaldehyde, permeabilized in saponin, and stained with Proteostat, diluted to 1:5000 in PBS, for 30 minutes at room temperature to detect protein aggregates. Finally, some cells were fixed in 1% paraformaldehyde and permeabilized in saponin before staining with 1:500 anti-pan-ubiquitin antibody (clone FK2; Sigma) followed by 1:500 goat anti-mouse IgG AF-555 secondary antibody (Thermo) to identify poly-ubiquitinated proteins. Cells were then washed twice in serum-free HBSS, resuspended in 300 ul of HBSS and analyzed using a FACS Lyric cytometer.

### Proteomic analysis

3×10^4^ LSK cells were double sorted and lysed in the buffer used for western blotting. Protein quantity was normalized to 10 µg per sample via Bicinchoninic Acid Assay (BCA). 35 µL of 5% SDS was added to each sample to bring the total volume to 50 µL. Following disulfide bond reduction and alkylation with tris(2-carboxyethyl)phosphine (Sigma-Aldrich, 646547-10x1ML) and iodoacetamide (Sigma-Aldrich, I1149-5G), samples were digested overnight with 1 µg of trypsin using S-Trap micro columns (Protifi). The peptide eluates from the S-Traps were dried and reconstituted in 50 µL of triethylammonium bicarbonate (TEAB, T7408-100ML) buffer. The TMT 6plex Isobaric Mass Tagging Kit (Thermo, 90066) was used to label the samples per the manufacturer’s instructions, quenched with 5% hydroxylamine, and the samples were combined. The samples were dried in a SpeedVac and reconstituted in 2% acetonitrile, 0.1% trifluoracetic acid buffer and diluted such that ∼1 µg of peptides were injected for each fraction.

Peptides were analyzed on a Thermo Orbitrap Eclipse MS system coupled to Ultimate 3000 RSLC-Nano liquid chromatography. Samples were injected onto a 75 µm internal diameter, 75-cm long EasySpray column (Thermo, ES905) and eluted with a gradient from 0-28% buffer B over 180 min at a flow rate of 250 nL/min. Buffer A contained 2% (v/v) ACN and 0.1% formic acid in water, and buffer B contained 80% (v/v) ACN, 10% (v/v) trifluoroethanol, and 0.1% formic acid in water. Spectra were continuously acquired in a data-dependent manner throughout the gradient, acquiring a full scan in the Orbitrap (at 120,000 resolution with a standard AGC target) followed by MS/MS scans on the most abundant ions in 2.5 s in the ion trap (turbo scan type with an intensity threshold of 5,000, CID collision energy of 35%, standard AGC target, maximum injection time of 35 ms and isolation width of 0.7 m/z). Charge states from 2-6 were included. Dynamic exclusion was enabled with a repeat count of 1, an exclusion duration of 25 s and an exclusion mass width of ± 10 ppm. Real-time search was used for selection of peaks for SPS-MS3 analysis, with the search performed against the human reviewed protein database from UniProt. Up to 1 missed tryptic cleavage was allowed, with carbamidomethylation (+57.0215) of cysteine and TMT reagent (+229.1629) of lysine and peptide N-termini used as static modifications and oxidation (+15.9949) of methionine used as a variable modification. MS3 data were collected for up to 10 MS2 peaks, which were matched to fragments from the real-time peptide search identification, in the Orbitrap at a resolution of 30,000, HCD collision energy of 65% and a scan range of 100–500.

Protein identification and quantification were done using Proteome Discoverer v.3.0 SP1 (Thermo). Raw MS data files were analyzed against the mouse reviewed protein database from UniProt (downloaded December 31, 2025, 17219 entries). Comet and SequestHT with INFERYS rescoring were used, with carbamidomethylation (+57.0215) of cysteine and TMT reagent (+229.1629) of lysine and peptide N-termini used as static modifications and oxidation (+15.9949) of methionine used as a variable modification. The Reporter Ions Quantifier node within Proteome Discoverer was used for quantitation, using reporter ion intensities. Peptide-spectrum matches were validated using the Percolator node within Proteome Discoverer. The false-discovery rate (FDR) cutoff was 1% for all peptides and proteins.

For proteomic analysis, protein raw abundance values were normalized using the Relative Log Expression (RLE) method and data were log2-transformed and imputed using the half-minimum imputation method before statistical tests were performed using Generalized Linear Models (GLMs) followed by Wald tests for statistical significance and Benjamini-Hochberg’s False Discovery Rate (FDR) method for multiple comparisons adjustments. The analysis was performed using the Morrison laboratory’s Omics Data Analyzer (ODA) 2.2.9 tool^78^. Gene set over-representation analysis was performed on genes that encoded proteins that significantly differed between *GCLC*-deficient and control LSK cells (fold change > 1.5 and FDR < 0.05). Proteins detected in any sample were used as background for the enrichment analysis. Proteins’ UniProt IDs were converted to gene symbols, and the enrichment analysis was done using ODA 2.2.9 with clusterProfiler 4.16.0, the msigdbr 10.0.2 database, and R 4.5.0.

### HSC colony forming assay

Bone marrow cells were stained with antibodies against HSC cell surface markers as described above. HSCs were sorted, one cell per well, into 96 well plates containing 100 µl of methylcellulose medium (M3434, STEMCELL Technologies) supplemented with 50 ng/ml of thrombopoietin (Peprotech) in the presence or absence of antioxidants and ferroptosis inhibitors. Ten to 12 days after sorting, colonies were counted using a brightfield microscope.

### Annexin V staining

1×10^7^ bone marrow cells were stained with antibodies against cell surface markers and washed twice with serum-free HBSS before staining with Annexin V protein conjugated to FITC in Annexin V binding buffer (10 mM HEPES, 140 mM NaCl, 2.5 mM CaCl_2_, pH 7.4). Samples were incubated for 15 minutes at room temperature in the dark with gentle shaking to prevent settling. After the incubation was complete, cells were washed twice with Annexin V binding buffer and resuspended in Annexin V binding buffer plus propidium iodide. Annexin V fluorescence was detected in the FITC channel by flow cytometry using a FACS Lyric.

### Statistical methods

In each type of experiment, multiple mice were tested in multiple independent experiments performed on different days. Mice were allocated to experiments randomly and samples processed in an arbitrary order, but formal randomization techniques were not used. No formal blinding was applied when performing the experiments or analyzing the data. Samples sizes were not pre-determined based on statistical power calculations but were based on our experience with these assays. No data were excluded.

Prior to analyzing the statistical significance of differences among treatments, we tested whether data were normally distributed and whether variance was similar among groups. To test for normality, we performed the Shapiro–Wilk tests when 3≤n<20 or D’Agostino Omnibus tests when n≥20. To test whether variability significantly differed among groups we performed *F*-tests (for experiments with two groups) or Levene’s median tests (for experiments with more than two groups). When the data significantly deviated from normality or variability significantly differed among groups, we log2-transformed the data and tested again for normality and variability. If the transformed data no longer significantly deviated from normality and equal variability, we performed parametric tests on the transformed data. If log2-transformation was not possible or the transformed data still significantly deviated from normality or equal variability, we performed non-parametric tests on the non-transformed data.

When data or log2-transformed data were normal and equally variable, statistical analyses were performed using Student’s *t*-tests (when there were two independent groups), one-way ANOVAs (when there were more than two independent groups), or matched samples two-way ANOVAs (when there were two or more groups with multiple cell populations or time points). When the data or log2-transformed data were normal but unequally variable, statistical analyses were performed using Welch’s *t*-tests (when there were two independent groups) or Welch’s one-way ANOVAs (when there were more than two independent groups). When the data and log2-transformed data were abnormal or unequally variable, statistical analysis was performed using Mann-Whitney tests (when there were two independent groups), Kruskal-Wallis tests (when there were more than two independent groups), or nparLD tests (when there were two or more groups measured at multiple time points). P-values from multiple comparisons were adjusted using Tukey’s (when there were more than two groups and all the comparisons were of interest) or Sidak’s method (when there were more than two groups and planned comparisons) after ANOVAs, Dunnett’s T3 method after Welch’s one-way ANOVAs, Dunn’s method after Kruskal-Wallis tests, or Benjamini-Hochberg’s False Discovery Rate (FDR) method after nparLD tests. Holm-Sidak’s method was used to adjust comparisons involving multiple cell populations with two groups. All statistical tests were two-sided where applicable. All data represent mean ± standard deviation. Statistical tests were performed using GraphPad Prism V11.1.0 or R 4.3.2.

## Acknowledgments

S.J.M. is a Howard Hughes Medical Institute Investigator, the Mary McDermott Cook Chair in Pediatric Genetics, the Kathryn and Gene Bishop Distinguished Chair in Pediatric Research, the director of the Hamon Laboratory for Stem Cells and Cancer, and a Cancer Prevention and Research Institute of Texas Scholar. T.P.M. and the Children’s Research Institute Metabolism Facility are supported by an award from the Cancer Prevention Research Institute of Texas (CPRIT Core Facilities Support Award RP24094). The BioHPC high-performance computing cloud at UTSW was used for data analysis and storage as well as proteomic data analysis (ODA) software deployment. We thank Michael Ortiz and Fernanda Lins and the Moody Foundation Flow Cytometry Facility for flow cytometry instrumentation and support, and Daniel Cassidy for help with mouse colony management. This article is subject to HHMI’s Open Access to Publications policy. HHMI lab heads have previously granted a nonexclusive CC BY 4.0 license to the public and a sublicensable license to HHMI in their research articles. Pursuant to those licenses, the author-accepted manuscript of this article can be made freely available under a CC BY 4.0 license immediately upon publication.

## Author Contributions

A.P. and S.J.M. conceived the project, planned the experiments, interpreted the results, and wrote the manuscript. A.P. performed all of the experiments with technical assistance from H.T., J.D.P., and S.M.. H.T. and G.H. performed western blots, helped with flow cytometry analyses, and helped to analyze the results. A.L performed the proteomics and sample preparation. T.P.M developed some of the metabolomic analysis methods and analyzed glutathione and vitamin E levels. Z.Z. developed the software for proteomic data analysis and performed the statistical analyses of the data.

## Declaration of interests

The authors declare no competing interests.

## Supplementary Figure Legends

**Supplementary Figure 1.**
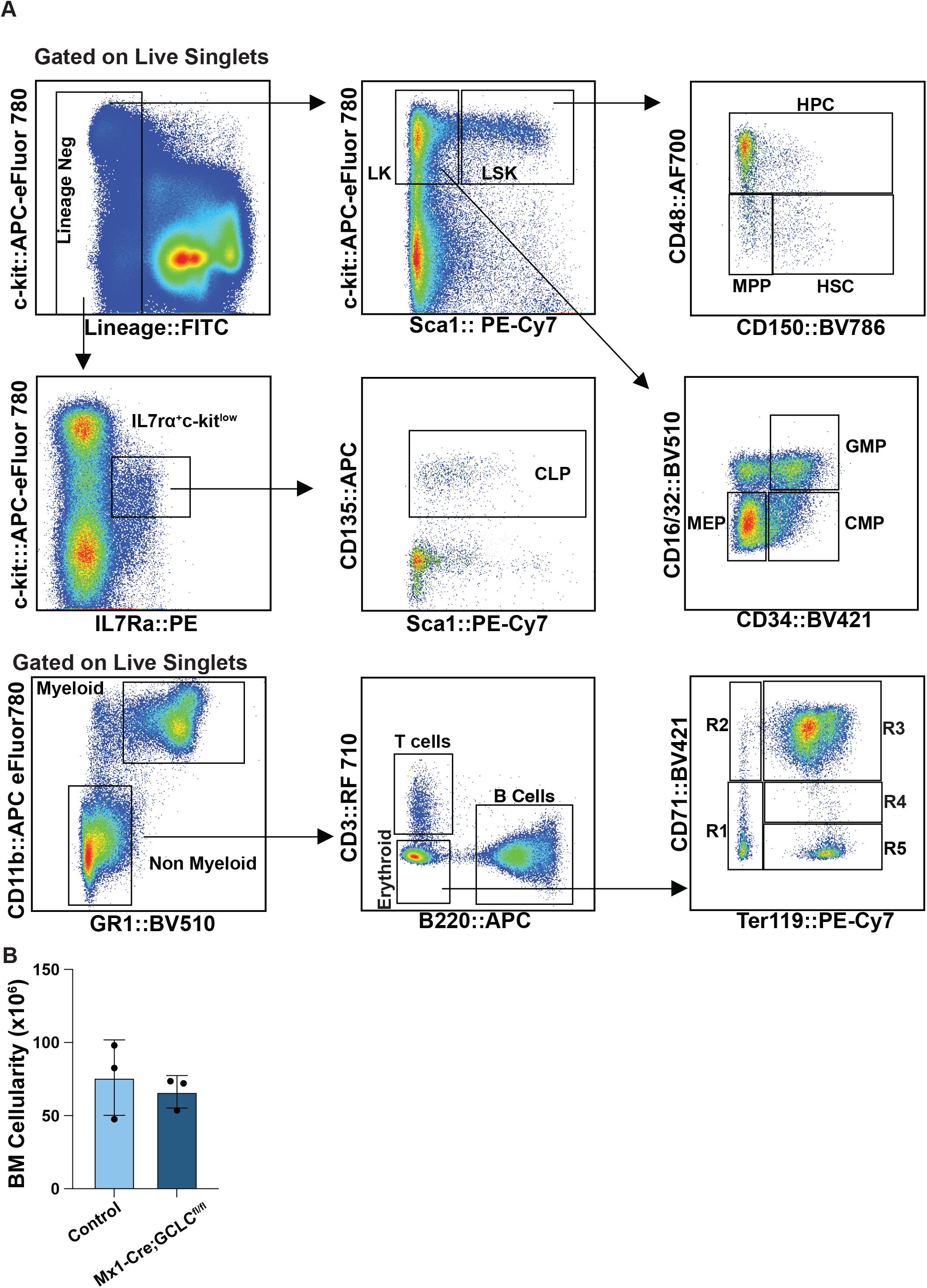
Flow cytometry gating strategy for the hematopoietic stem and progenitor cell populations characterized in this study. (A) Flow cytometry plots detailing the markers and gating strategies used to analyze the hematopoietic stem and progenitor cell populations characterized in this study. (B) Bone marrow cellularity in recipient mice transplanted with equal numbers of *Mx1-cre; GCLC^fl/fl^* and control bone marrow cells, then treated with poly I:C 6 weeks later and analyzed 8 weeks after poly I:C treatment. Mice reconstituted by a mixture of *GCLC*-deficient and control cells did not develop pancytopenia (3 recipient mice per genotype). Each dot represents a different mouse and data represent mean ± standard deviation (**b**). Statistical significance was assessed using a two-sided Student’s *t*-test (**b)**.

**Supplementary Figure 2.**
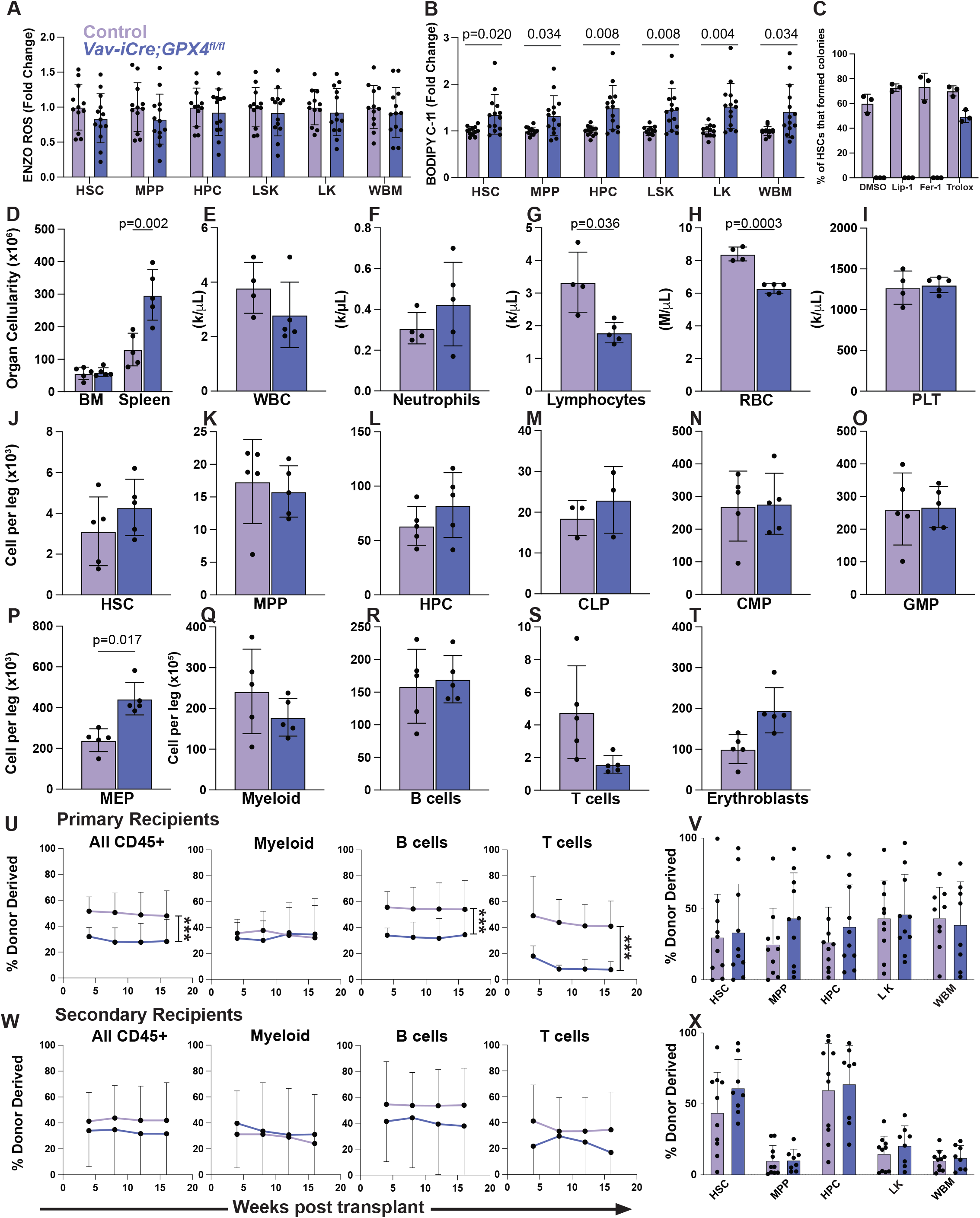
Deletion of *GPX4* from hematopoietic cells had little effect on hematopoiesis or HSC function. (A-B) Fold change in median fluorescence intensity of ENZO Total ROS (A) or BODIPY C-11 (B) staining of HSCs, MPPs, HPCs, LSKs, LK, and whole bone marrow cells from *Vav-iCre;GPX4^fl/fl^* versus control mice (a total of 12-14 mice per genotype from 4 independent experiments). (C) Colony formation by HSCs from *Vav-iCre;GPX4^fl/fl^* or control mice cultured in DMSO (vehicle), 1uM Liprostatin-1, 1uM Ferrostatin-1, or 300 uM Trolox (a total of 3 replicates per treatment from 3 independent experiments). (D-I) Bone marrow and spleen cellularity (D) as well as white blood cell (E), neutrophil (F), lymphocyte (G), red blood cell (H), and platelet (I) counts from *Vav-icre;GPX4^fl/fl^* and littermate control mice (a total of 4-5 mice per genotype from two independent experiments). (J-T) Numbers of HSCs (J), MPPs (K), HPCs (L), CLPs (M), CMPs (N), GMPs (O), MEPs (P), Mac1^+^CD11b^+^ Myeloid cells (Q), B220^+^ B cells (R), CD3^+^ T cells (S), and CD71^+^Ter119^+^ basophilic erythroblasts (T) in the bone marrow (one femur and tibia) of *Vav1-iCre;GPX4^fl/fl^* and littermate control mice (a total of 5 mice per genotype in two independent experiments). (U-V) Donor cell contributions to the blood (U) and bone marrow (V) of mice competitively transplanted with *Vav-icre; GPX4^fl/fl^* or control donor bone marrow (2 donors per genotype were transplanted into 8-10 recipient mice per genotype in 2 independent experiments). (W-X) Donor cell contributions to the blood (W) and bone marrow (X) of secondary recipient mice transplanted with whole bone marrow cells from the primary recipient mice analyzed in Suppl. Figure 2U (2-3 primary recipient mice per genotype were transplanted into 8-10 secondary recipient mice per genotype from two independent experiments). All dots represent a different mouse except in panels U and W. All data represent mean ± standard deviation. Statistical significance was assessed using matched samples two-way ANOVAs followed by Sidak’s multiple comparisons adjustments (**a,d**), Mann-Whitney tests followed by Holm-Sidak’s multiple comparisons adjustments (**b,c**), Student’s *t*-tests followed by Holm-Sidak’s multiple comparisons adjustments (**e-t,v,x**), or nparLDs followed by the FDR multiple comparisons adjustments (**u,w**). All statistical tests were two-sided.

**Supplementary Figure 3.**
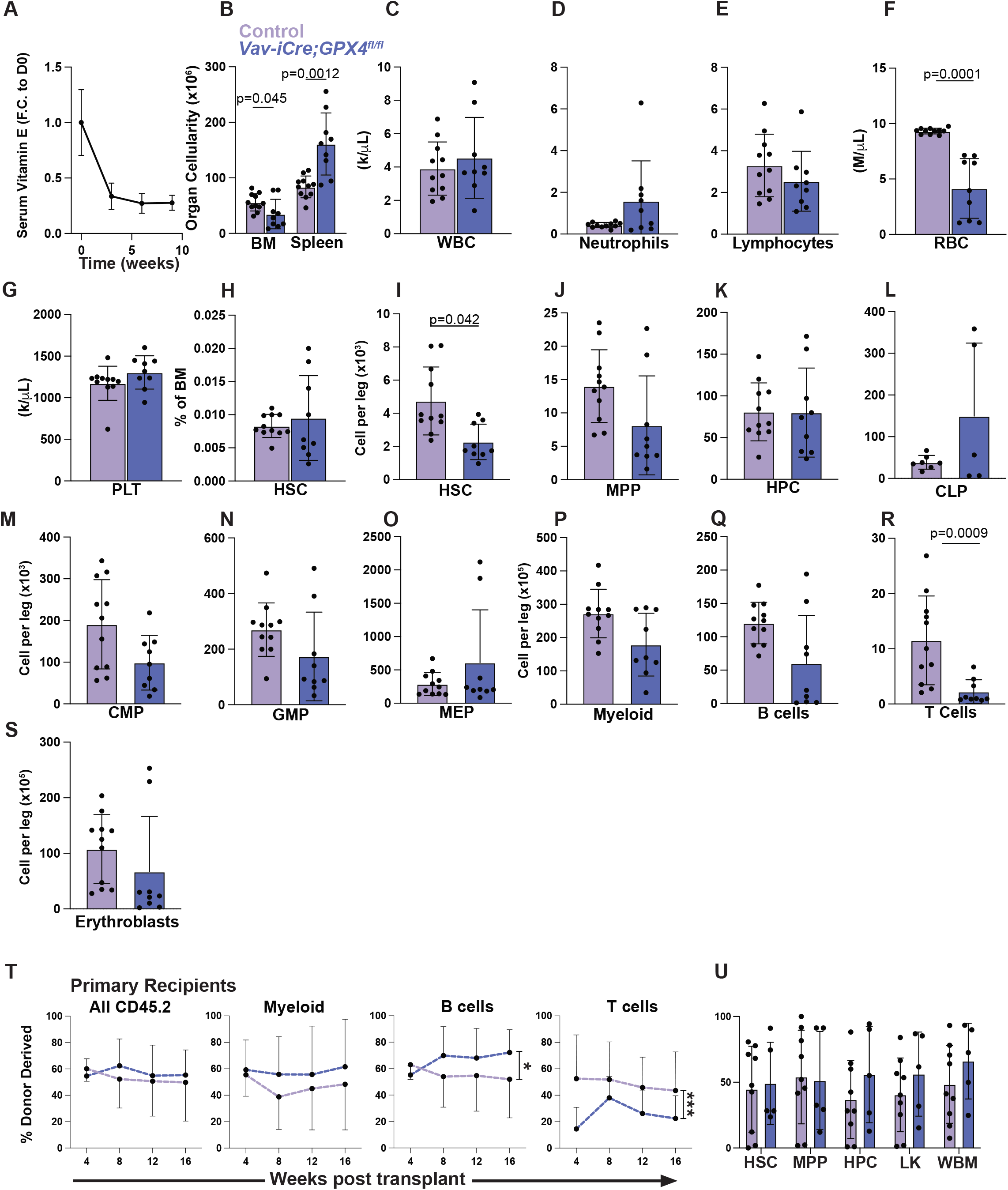
Even after vitamin E depletion, *GPX4* deficiency did not affect HSC function. (A) Relative serum vitamin E levels over time in wild-type mice fed a low vitamin E diet (a total of 5 mice per time point in two experiments). (B-G) Bone marrow and spleen cellularity (B) as well as white blood cell (C), neutrophil (D), lymphocyte (E), red blood cell (F), and platelet (G) counts in *Vav-icre; GPX4^fl/fl^* and littermate control mice fed a low vitamin E diet for three weeks prior to analysis (a total of 9-11 mice per genotype from three independent experiments). (H, I) Frequency (H) and number (I) of HSCs in the bone marrow (one femur and tibia) of *Vav-icre; GPX4^fl/fl^* and control mice fed a low vitamin E diet for three weeks (a total of 9-11 mice per genotype in three independent experiments). (J-S) Numbers of MPPs (J), HPCs (K), CLPs (L), CMPs (M), GMPs (N), MEPs (O), Mac1^+^CD11b^+^ Myeloid cells (P), B220^+^ B cells (Q), CD3^+^ T cells (R), and CD71^+^Ter119^+^ basophilic erythroblasts (S) in the bone marrow (one femur and tibia) of *Vav-icre; GPX4^fl/fl^* and littermate control mice fed a low vitamin E diet for three weeks (a total of 9-11 mice per genotype in three independent experiments). (T-U) Donor cell contributions to the blood (T) and bone marrow (U) of mice competitively transplanted with *Vav-icre; GPX4^fl/fl^* or control donor bone marrow from mice fed a low vitamin E diet for three weeks. The recipient mice were also maintained on a low vitamin E diet for the duration of the experiment, starting three weeks before transplantation (2 donors per genotype were transplanted into 5-9 recipient mice per genotype in 2 independent experiments). Each dot represents a different mouse except in panels A and T. All data represent mean ± standard deviation. Statistical significance was assessed using Mann-Whitney tests followed by Holm-Sidak’s multiple comparison’s adjustments (**b-g** and **i-s**), a Mann-Whitney test (**h**), nparLD followed by FDR multiple comparisons adjustment (**t**), or Student’s *t*-tests followed by Holm-Sidak’s multiple comparisons adjustments (**u**). All statistical tests were two-sided.

**Supplementary Figure 4.**
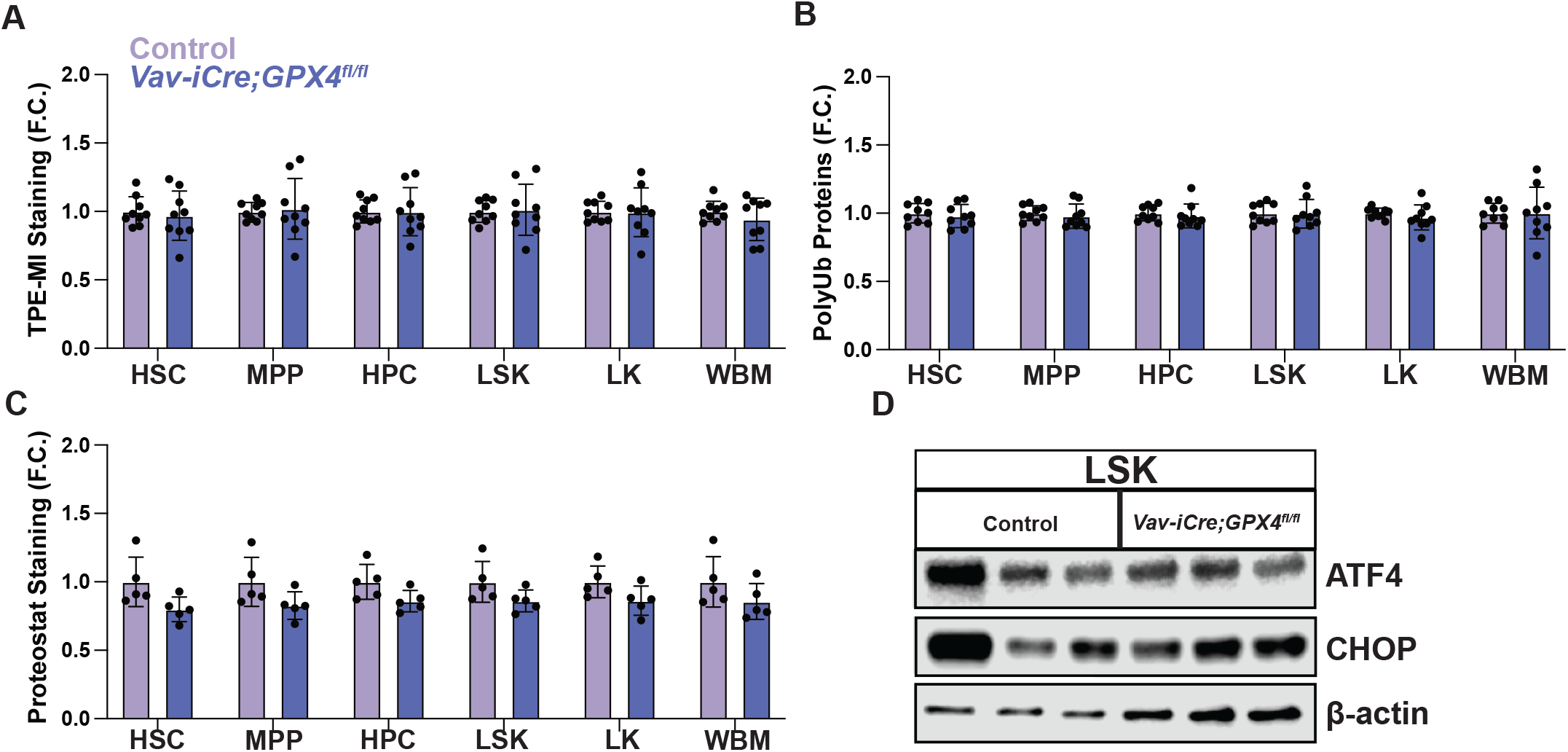
*GPX4* deletion did not induce an unfolded protein response in hematopoietic cells. (A) Fold change in median fluorescence intensity of tetraphenylethene malemide fluorescence (TPE-MI) in HSCs, MPPs, HPCs, LSKs, LK, and whole bone marrow cells in *Vav-icre; GPX4^fl/fl^* versus littermate control mice. Fold change is relative to the respective control cell type. (n=9 mice per genotype from two independent experiments) (B) Fold change in median fluorescence intensity of staining for polyubiquitinated proteins in HSCs, MPPs, HPCs, LSKs, LK, and whole bone marrow cells from *Vav-icre; GPX4^fl/fl^* and littermate control mice, normalized to whole bone marrow cells in each experiment (a total of 9 mice per genotype from two independent experiments). (C) Fold change in median fluorescence intensity of Proteostat staining in HSCs, MPPs, HPCs, LSKs, LKs, and whole bone marrow cells from *Vav-icre; GPX4^fl/fl^* versus control mice (a total of 5 mice per genotype from two independent experiments). (D) Western blot analysis of ATF4 and CHOP levels relative to β-actin loading control in 100,000 LSK cells per lane from *Vav-icre; GPX4^fl/fl^* and littermate control mice (one mouse per lane). Each dot represents a different mouse. All data represent mean ± standard deviation. Statistical significance was assessed using Mann-Whitney tests followed by Holm-Sidak’s multiple comparisons adjustments (**a,b**), or Student’s *t*-tests followed by Holm-Sidak’s multiple comparisons adjustments (**c**). All statistical tests were two-sided.

## STAR Methods

The key resources table is uploaded as a separate document.

## Lead Contact

Further information and requests for resources and reagents should be directed and will be fulfilled by the lead contact, Sean J. Morrison.

## Data and Code Availability

- All data reported in this paper will be shared by the lead contact upon reasonable request.
- This paper does not report original code.
- Mass spectrometry data used for proteomic analysis were deposited in the MassIVE database (accession number: MSV000103217) and the proteomics data themselves were deposited in the ProteomXchange database (accession number: PXD084161).

---

**Table S1:**
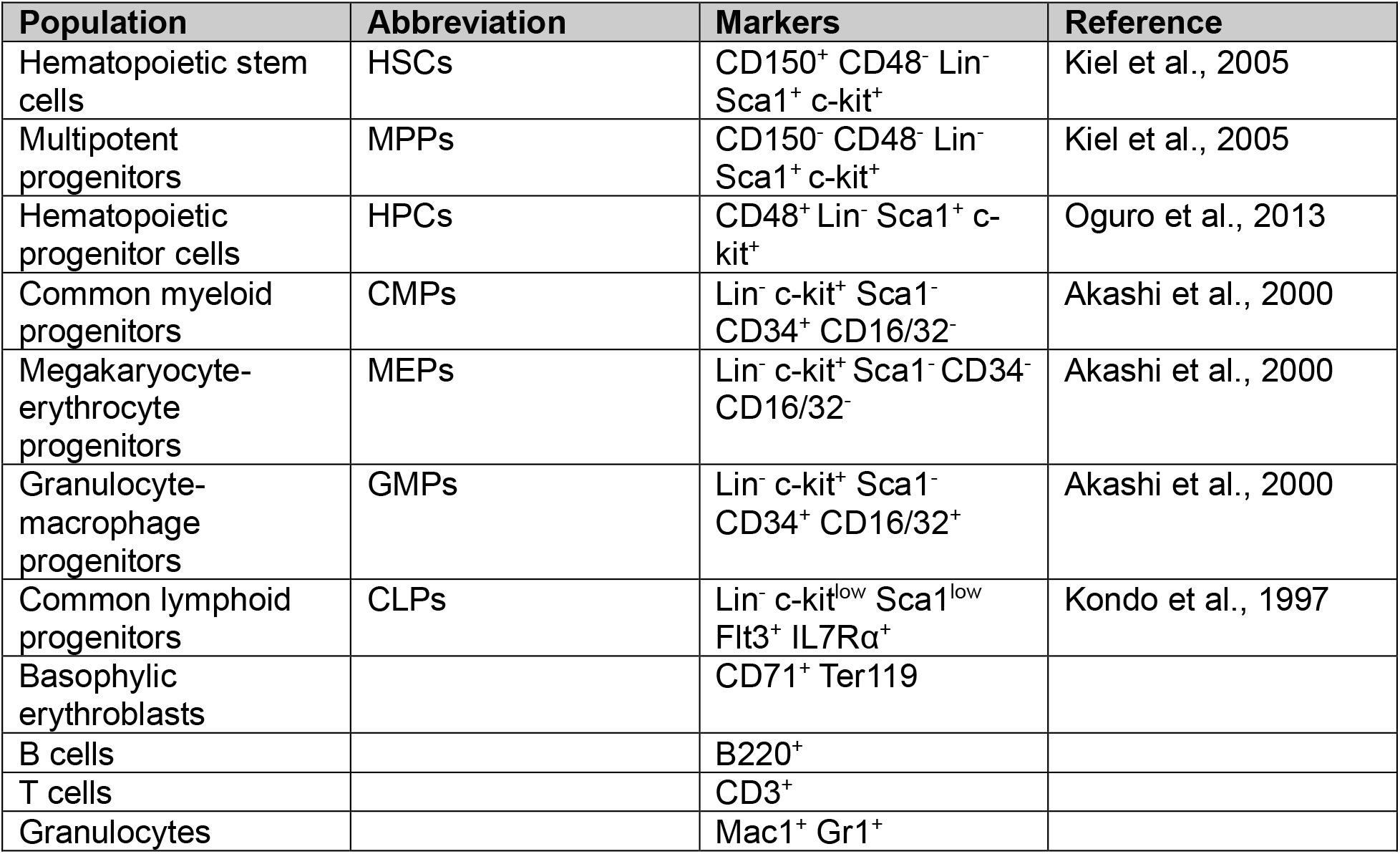
Related to Figure 1 and Supplementary Figure 1. Cell populations analyzed by flow cytometry in this study.

**Table S2.**
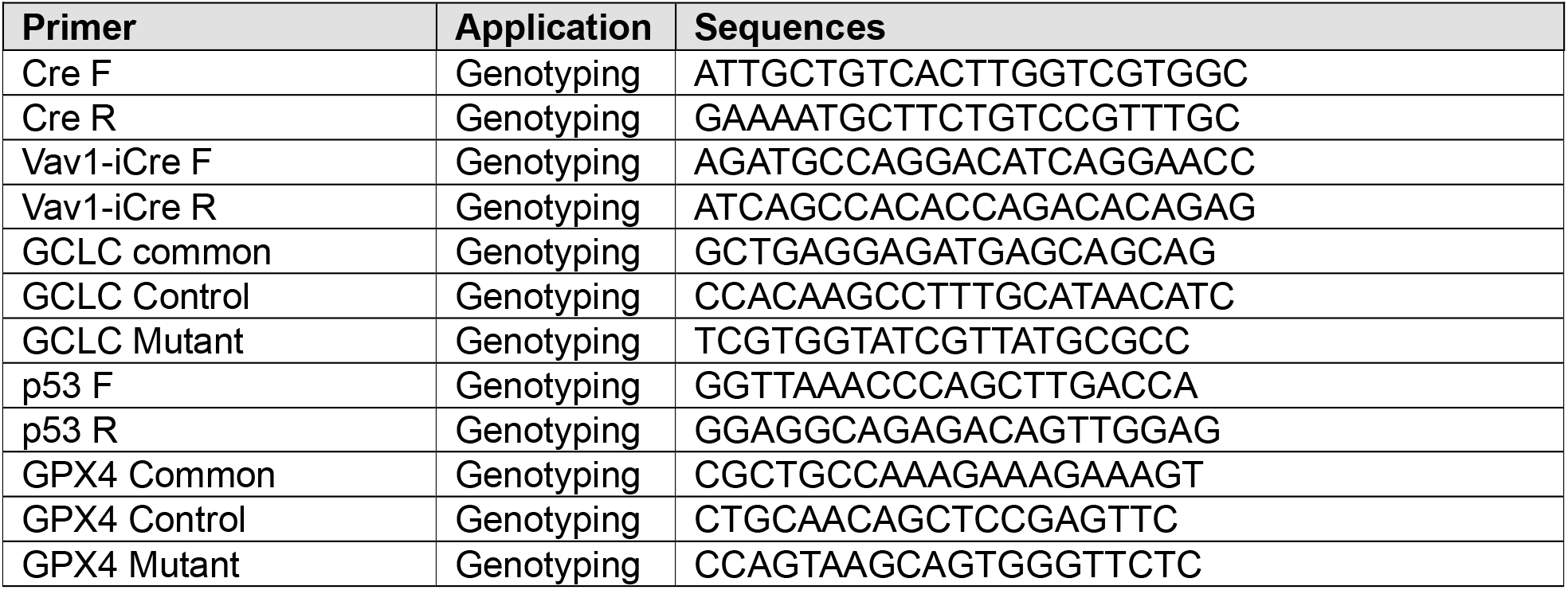
Related to STAR Methods. Primers used in this study.

## KEY RESOURCES TABLE

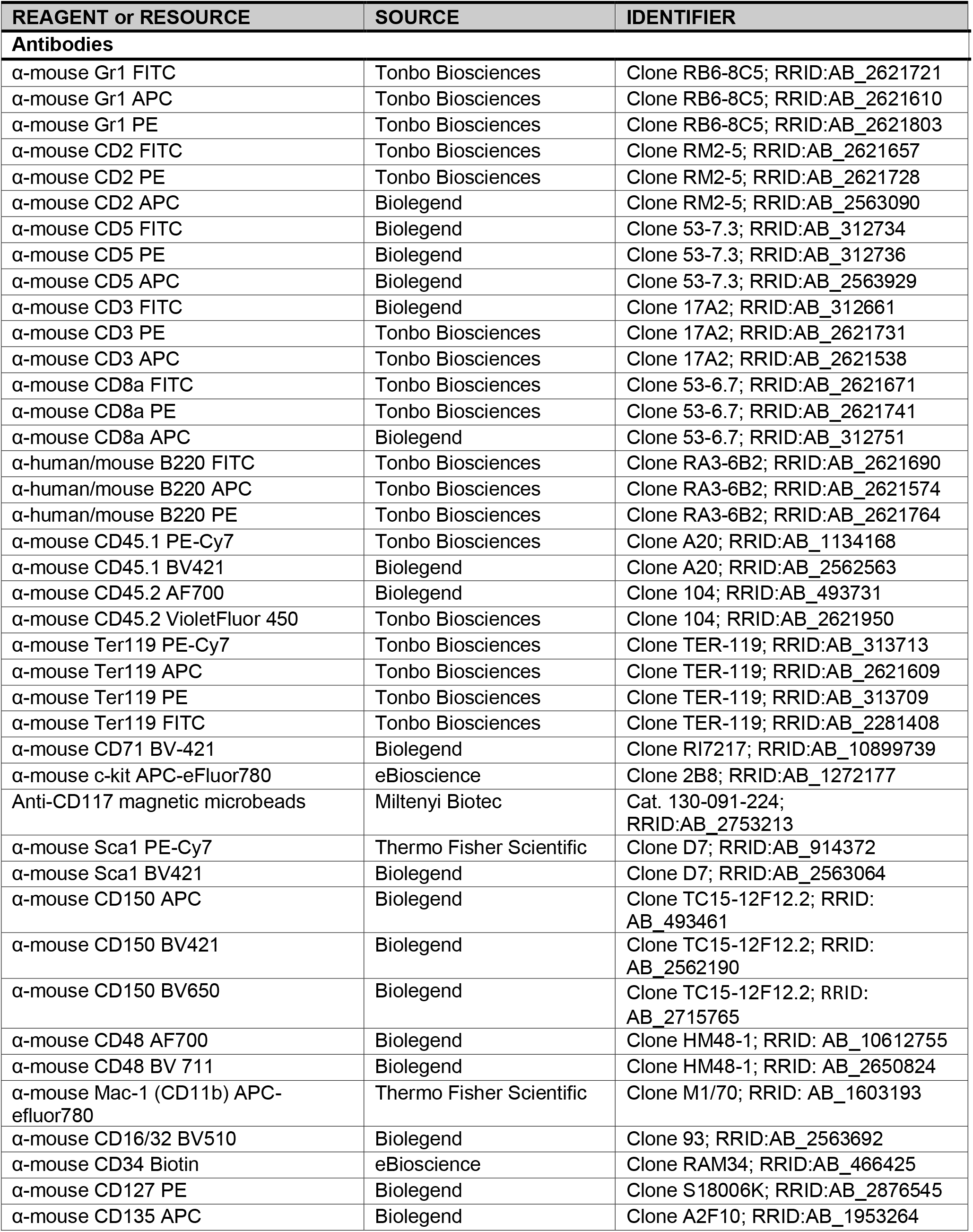
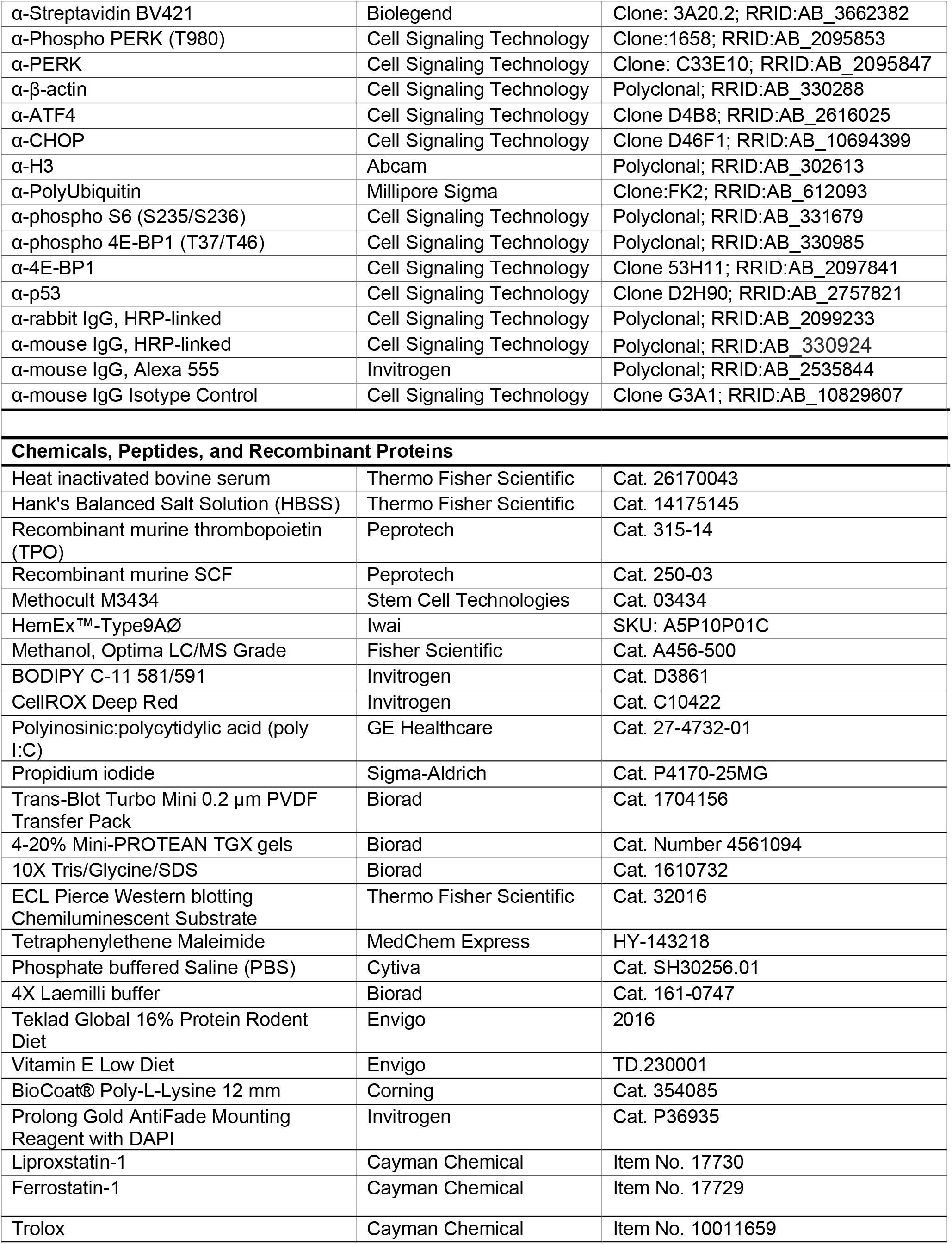
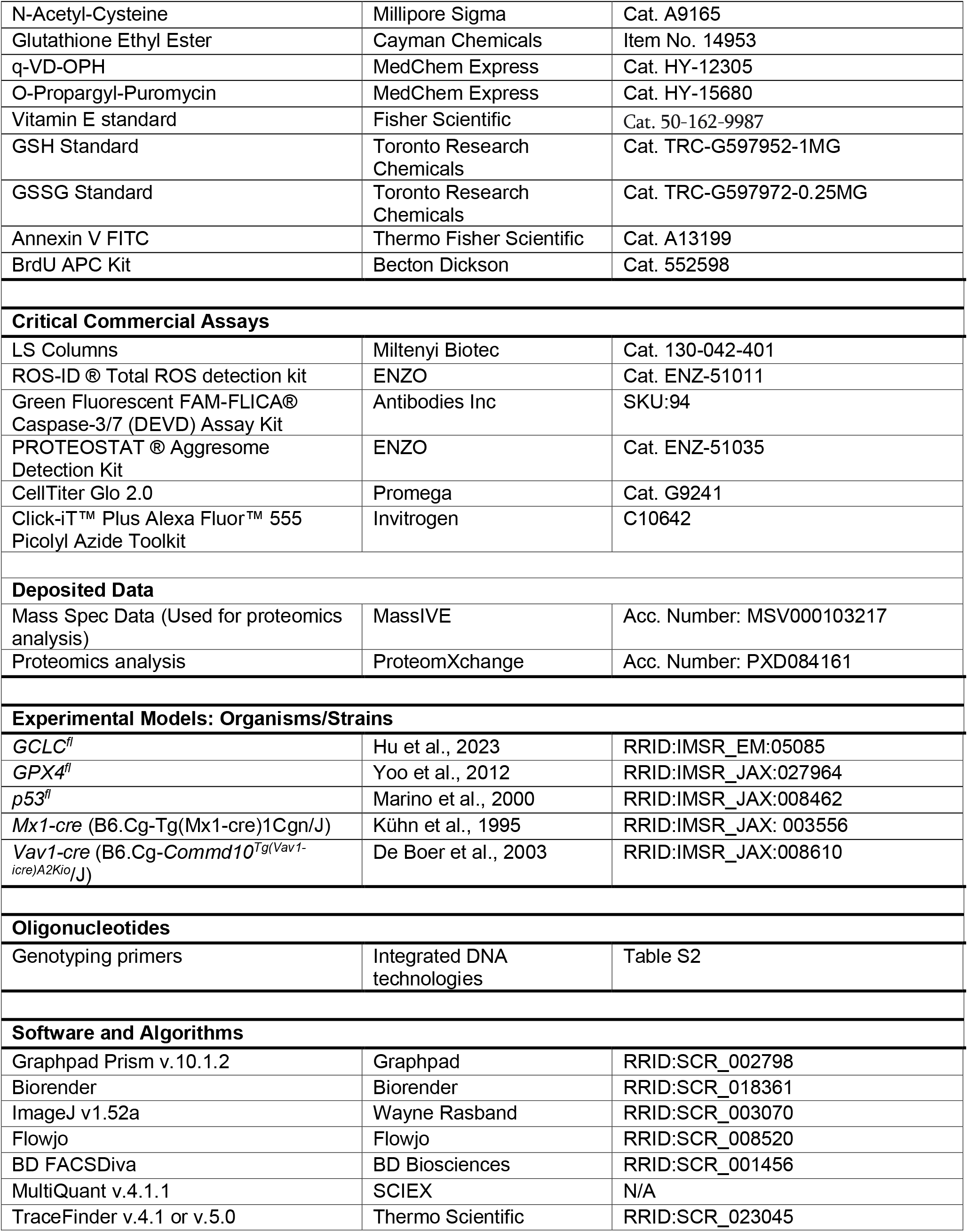

## Notes

### Competing Interest Statement

The authors have declared no competing interest.

